# Position-dependent Codon Usage Bias in the Human Transcriptome

**DOI:** 10.1101/2021.08.11.456006

**Authors:** Kaavya Subramanian, Nathan Waugh, Cole Shanks, David A Hendrix

**Author notes:** these authors contributed equally to this work (Kaavya Subramanian, Nathan Waugh).

## Abstract

All life depends on the reliable translation of RNA to protein according to complex interactions between translation machinery and RNA sequence features. While ribosomal occupancy and codon frequencies vary across coding regions, well-established metrics for computing coding potential of RNA do not capture such positional dependence. Here, we investigate *position-dependent codon usage bias* (*PDCUB*), which dynamically accounts for the position of proteincoding signals embedded within coding regions. We demonstrate the existence of PDCUB in the human transcriptome, and show that it can be used to predict translation-initiating codons with greater accuracy than other models. We further show that observed PDCUB is not accounted for by other common metrics, including position-dependent GC content, consensus sequences, and the presence of signal peptides in the translation product. More importantly, PDCUB defines a spectrum of translational efficiency supported by ribosomal occupancy and tRNA adaptation index (tAI). High PDCUB scores correspond to a tAI-defined translational ramp and low ribosomal occupancy, while low PDCUB scores exhibit a translational valley and the highest ribosomal occupancy. Finally, we examine the relationship between PDCUB intensity and functional enrichment. We find that transcripts with start codons showing the highest PDCUB are enriched for functions relating to neuropeptide signaling and nucleosome assembly, as well as development of musculoskeletal, cardiovascular, neurological, gastrointestinal, sensory, and other body systems. Furthermore, transcripts with high PDCUB are depleted for functions related to immune response and detection of chemical stimulus. These findings lay important groundwork for advances in our understanding of the regulation of translation, the calculation of coding potential, and the classification of RNA transcripts.

## INTRODUCTION

Translation from RNA to protein is a fundamental and ubiquitous life process. Some studies estimate that ≈68% of human genes do not encode proteins, and are transcribed as long noncoding RNAs (lncRNAs) or transcripts of unknown coding potential [1]. This raises the question of how the ribosome distinguishes mRNAs from lncRNAs that have open reading frames (ORFs) [2]. At the same time, other studies suggest that some lncRNAs are weakly translated [3–5]. These observations underscore the importance of understanding how the ribosome uses sequence features to distinguish mRNAs from lncRNAs, as well as to properly identify the start codon within an mRNA. Such sequence features could help explain the translation of short transcripts that encode small peptides [6, 7], and improve the design of mRNA vaccines [8].

The Kozak consensus sequence is one such motif, discovered by Marilyn Kozak in the 1980s [9]. This sequence characterizes and helps identify start codons in eukaryotic mRNAs and is regarded as a key feature that enhances translational potential. Another approach to quantifying mRNA coding potential is codon usage bias (CUB), which refers to the frequency of usage for each codon in the coding portion of a transcriptome relative to the frequency of synonymous codons. Computational models of CUB have been shown to perform well in predicting translational efficiency and also show an association with ribosome profiling data [10]. The codon adaptation index (CAI) is a widely recognized metric of CUB that assigns a score to each transcript based on its length and codon composition relative to overall CUB across the coding transcriptome [11]. The CAI model has been used as a baseline for other quantitative models of CUB, and has also been correlated with gene expression in select genomes [11].

A potential limitation of these approaches is that they do not consider the role of codon position or transcript length explicitly. Recently, studies of position-dependent codon usage bias (PDCUB) observed that codon usage is non-uniform with regard to transcript position in *E. coli* [12] and yeast [13]. Moreover, our own investigations demonstrated that a recurrent neural network, which we trained to distinguish mRNAs from lncRNAs based on sequence alone, was able to learn sequence-specific rules and make classification decisions approximately 100-200 nucleotides (nt) downstream of the start codon [14].

Here, we investigate the extent of PDCUB within the human coding transcriptome and whether PDCUB can function as a predictor of start codon location. We present a codon bias score that incorporates these position-dependent codon patterns, using only information observed in the first 300 nt downstream of a given AUG. We show that our score is capable of distinguishing start codons from non-start AUGs with greater accuracy than other methods. While our PDCUB score was constructed from general positional trends in codon usage, we show that it is correlated with translational efficiency data. We show that the highest PDCUB transcripts are associated with a translational ramp and low ribosomal occupancy, while the lowest are associated with a translational valley and high ribosomal occupancy. Moreover, we show that high- and low-PDCUB transcripts display a clear division in biological function, respectively corresponding to a need for high and low translational efficiency.

## RESULTS

### Visualization of PDCUB

We first performed a coarse-grained investigation of PDCUB in the human transcriptome using GENCODE data [15]. The frequencies of all 61 sense codons in protein-coding transcripts were visualized in successive 60-nt bins, up to 3000 nt downstream of the annotated start codon. We transformed the observed bin-specific codon frequencies to z-scores, defined relative to the global average frequency of codon occurrences within a bin, across the 50 bins downstream of the AUG. We observed that most codons displayed a distinct position-dependent z-score profile as we move downstream from the start codon, with some codons increasing in usage and others decreasing (Figure 1A). This led us to evaluate a more refined model of PDCUB in the human transcriptome.

**Figure 1:**
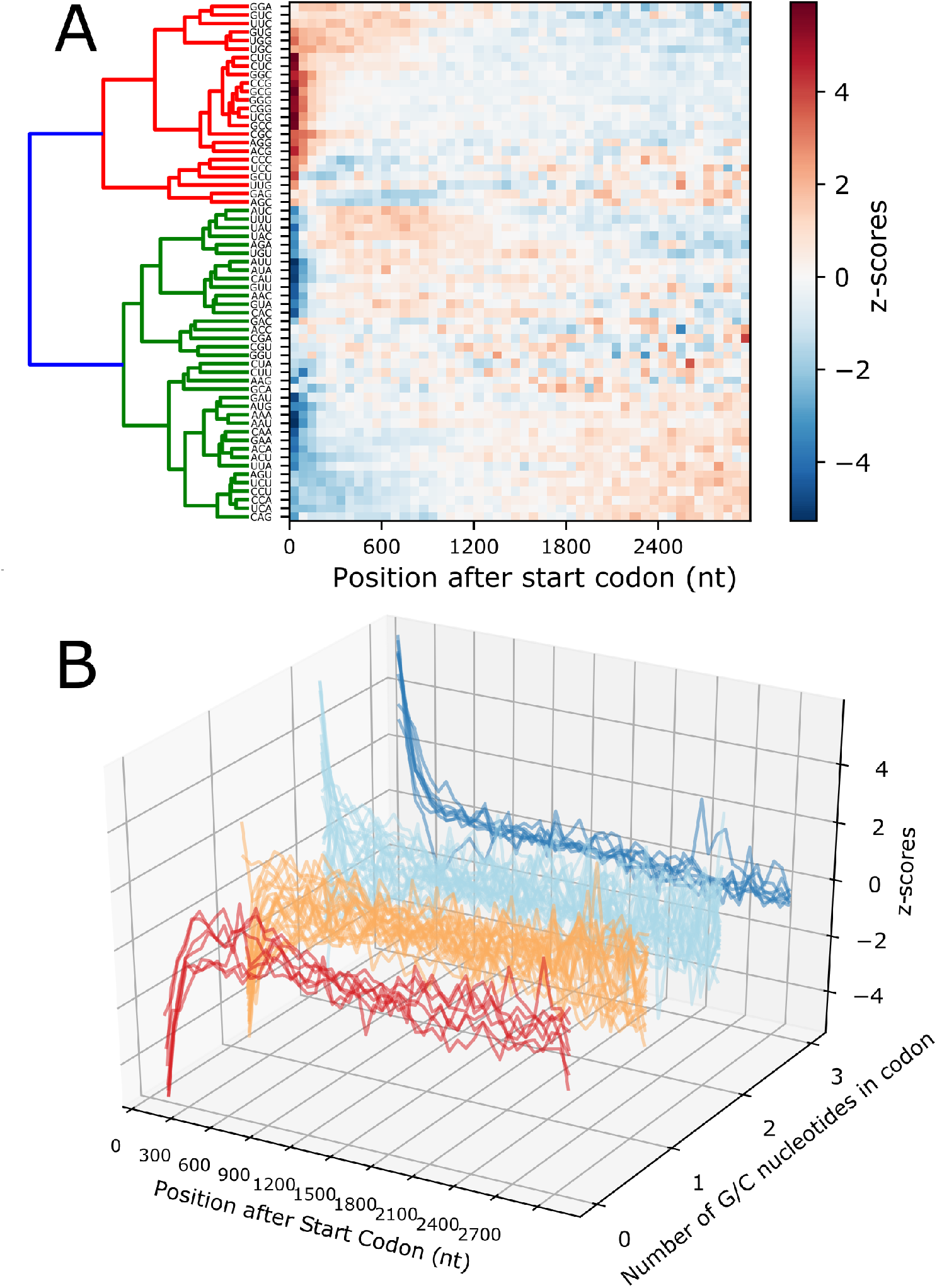
**A.** A “bird’s eye view” of position-dependent codon usage bias in the human transcriptome (GENCODE protein-coding transcript models). A heatmap and dendrogram visualizing the z-scores computed from codon occurrence frequency in bins of size 60nt. **B.** Curves show the z-scores computed from codon occurrence frequencies, shown in clusters corresponding to the number of G/C nucleotides within the codon.

We investigated whether the heatmap of z-score profiles reveals other patterns in position-dependent codon usage when the codons are clustered according to distinct criteria. First, we organized codons by their amino acid chemical properties, but did not observe a strong correspondence between z-score profile and encoded amino acid chemical type (Supplementary Figure 1). We next performed hierarchical clustering on the codons and found that they were predominantly organized by G and C (GC) nucleotide content. To test this observation, we compared our computed z-score dendrogram distance against a modified Hamming distance that compares the extent of variation of GC nucleotides between two codons (Figure 1A). We found a strong correspondence between the number of common GC nucleotides and the dendrogram distance in Figure 1A, suggesting that the GC content of each codon is indeed a significant factor in the pattern of position-dependent z-scores (Supplementary Figure 2).

Next, we grouped codons by their GC content and plotted z-score curves for each individual codon (Figure 1B). We found that codons consisting exclusively of GC nucleotides show a strong enrichment near the start codon, while codons consisting exclusively of A or U (AU) nucleotides are depleted immediately after the start codon. Codons with either 1 or 2 GC nucleotides continue this trend, with GC-rich codons (2 GC nucleotides) showing greater similarity to GC-only codons, while GC-poor codons (1 GC nucleotide) show greater similarity to AU-only codons.

### Optimizing PDCUB weight matrix

We used the prediction of start codons to assess the importance of the observed PDCUB patterns in the human transcriptome. We created training, validation, and test sets by subdividing the protein-coding transcripts from GENCODE using an 80:10:10 split. We created PDCUB position-specific scoring matrices (PSSMs, or weight matrices) to score the occurrence of a codon *c* at bin *b* of a potential open reading frame, relative to the global frequency of the nucleotide triplet corresponding to the codon *c*. We assigned the PDCUB weight matrix score (PDCUB score) to the series of triplets that follow each AUG in the transcripts across all frames, to discern whether the PDCUB score was sufficient to distinguish start codons from all other AUGs in a transcript.

We focused on the first 300 nt downstream of the start codon so that our score could be applied to most mRNA and lncRNA transcripts, including those with shorter ORFs. As a metric of predictive performance, we computed a recall, defined as the percentage of transcripts in which our PDCUB score identified the start codon as the highest-scoring AUG in that transcript. We observed that start codon recall depends on bin number and bin size, so we systematically computed the recall for combinations of both parameters (Figure 2A). We computed our PDCUB weight matrix using the training set using different combinations of bin number and size, selected an optimal set of parameters using performance on the validation set, and computed final percentages on the test set. We found that start codons were detected with maximum accuracy using one codon per bin – in other words, the combination of 100 single-codon bins provided the greatest predictive accuracy (Figure 2A). The z-score heatmap for this optimal bin size and number of bins is presented in Figure 2B. Hierarchical clustering of this heat map also corresponds to codon GC-content.

**Figure 2:**
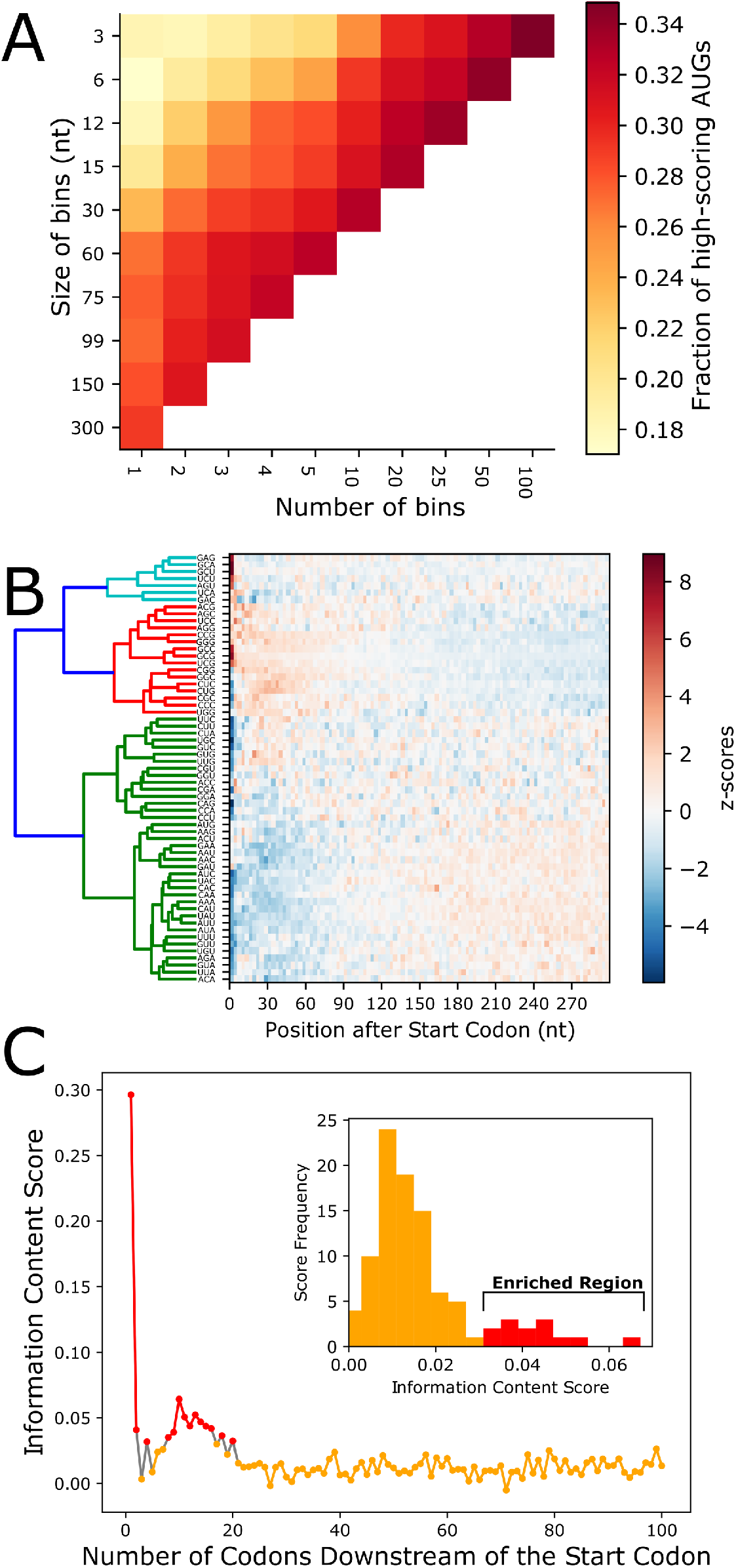
**A.** A heatmap demonstrating the optimal bin size and number of bins within the first 300 nt of CDS regions of the human transcriptome favors 100 3-nt bins. **B.** A codon z-score heatmap weight hierarchical clustering for the 300 nt after the start codon **C.** The information content of all transcripts, computed from position-dependent codon frequencies shows a region of enriched information content relative to the background trinucleotide frequencies. Red line segments indicate some regions of enriched information content. The inset shows a histogram of the information content for the first 100 codons with a tail of enriched information content, shown in red.

### Information content of the PDCUB model

We used the PDCUB probability matrix to compute the information content as a function of nucleotide position or bin (see Methods). We observed enriched information content in the range from codons 0 to 20, relative to and downstream of the start codon (Figure 2C).

### Start codon identification as a function of transcript length

We next investigated the role of transcript length in the percentage of correctly identified start codons – specifically, whether transcript length is correlated with the percentage of start codons identified by our PDCUB score as the highest-scoring AUG triplet in a given transcript. We compared the central tendencies (mean and median) for PDCUB scores computed for start codons and for other AUGs. We found that there was a significant difference between start codons and non-start AUGs for all transcript lengths (Supplementary Figure 3).

We compared the length-dependent recall of the PDCUB score with four other predictive models. First, we compared the PDCUB model to a Kozak consensus sequence model, which relies on finding within a transcript the nucleotide sequence of fewest possible mismatches to the Kozak consensus sequence [16]. We also compared the PDCUB model to a weight matrix computed from base composition in the region of the Kozak sequence, from −10 to +1 relative to the start codon. A null model that estimates the rate for the random selection of start codons amongst all AUG triplets, and a position-dependent GC-content model, were also used as controls. We found that the PDCUB model consistently performed best among the five models (Figure 3A). Even though PDCUB only considers the first 300 nt of the coding sequence, we observed improved performance with shorter transcripts. Figure 3B shows an example transcript, plotting the PDCUB score over all positions, and plotted separately for each reading frame. This example transcript conforms to the trend observed in the majority of transcripts of this length, with the start codon having the highest PDCUB score.

**Figure 3.**
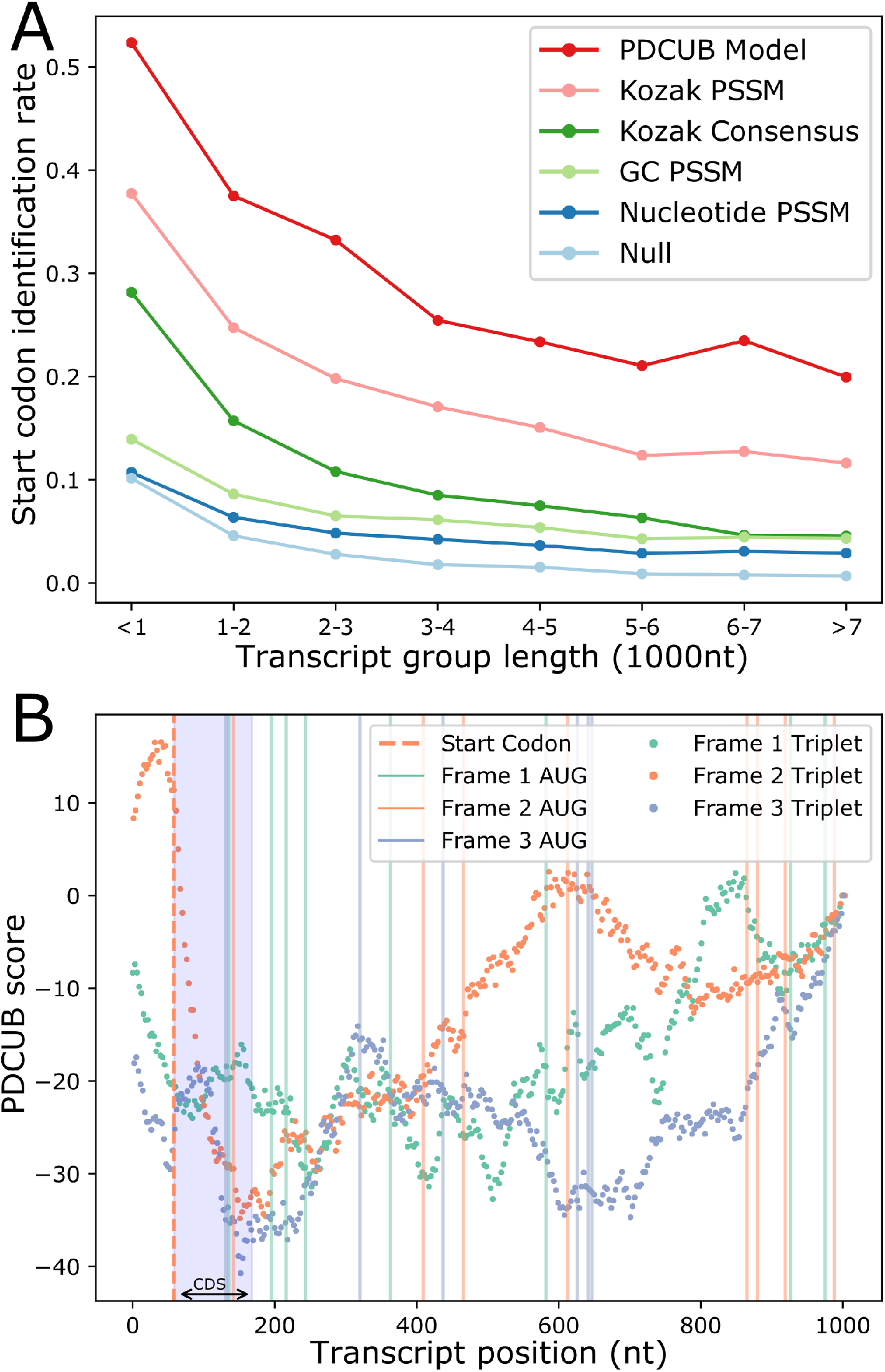
**A.** The rate of start codon identification compared for different models. The curves show what fraction of start codons are identified per transcript, comparing the PDCUB score computed from a PSSM compared to the Kozak consensus sequence AUGGCC, a PSSM computed from the positions −10 to +2 relative to the start codon for all transcripts, a PSSM computed from the position-dependent GC content within CDS regions relative to global GC content, a PSSM computed from single-nucleotide frequencies in CDS regions relative to global nucleotide frequencies, a random selection model. **B.** The PDCUB score plotted as a function of position, starting at each codon, and plotted separately for each reading frame, for an example short (~1000nt) transcript. Blue shaded area beginning at the start codon indicates the CDS region.

### Comparison of mRNAs and lncRNAs with PDCUB scores

We compared scoring mRNAs using the annotated CDS to lncRNAs by scoring the longest ORF in each GENCODE lncRNA with an ORF using the PDCUB scores. We found that in general, PDCUB was substantially higher in mRNAs than in lncRNAs (Supplementary Figure 3). We performed a Kolmogorov-Smirnov test to evaluate the difference between the mRNA PDCUB distribution to the distribution for lncRNAs, and it resulted in a p-value of 0.731e-07. We further subdivided both the mRNAs and lncRNAs into different ranges of 1000 nt. Supplementary Figure 4 shows the mean and median values for each of these bins, along with 95% confidence intervals.

### Functional enrichment in high-PDCUB transcripts

We identified a subset of transcripts with particularly high-significance PDCUB scores relative to the distribution of scores observed over all AUG triplets (Supplementary Figure 4) and named this our significant PDCUB set. We also divided our total transcript set into quintiles based on PDCUB score (see methods). We examined whether transcripts in these sets were enriched for biological functions or protein sequence patterns that might explain their scores.

#### Gene Ontology (GO) enrichment

We investigated whether the transcripts in our high-significance and low-significance PDCUB transcript sets, as well as in the various PDCUB quintiles (see methods), were enriched or depleted for particular biological functions. A table of significant Gene Ontology (GO) terms is provided in Supplementary Table 1. A similar analysis was performed for CAI quintiles as a control (also in Supplementary Table 1, in different sheets). Enrichment of significant GO terms associated with transcripts in the bottom PDCUB quintile is visualized in a scatterplot (Figure 4A). Similarly, GO terms that are enriched in the top quintile are shown in Figure 4B.

**Figure 4:**
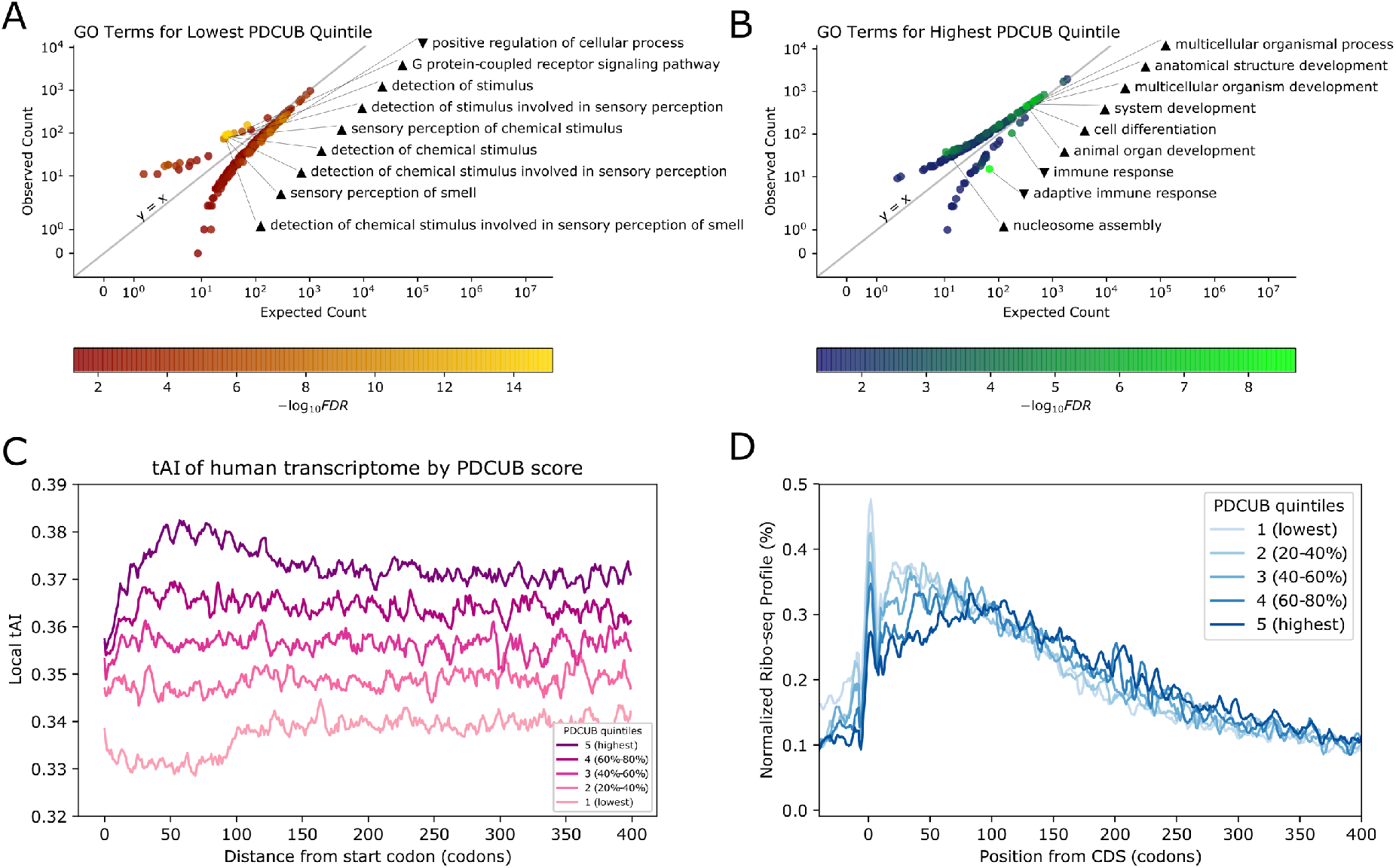
**A.** Scatterplot representing statistically enriched GO terms associated with transcripts in the lowest PDCUB score quintile. Each dot corresponds to an enriched GO term, the x-axis position corresponds to the expected number of occurrences, the y-axis represents the observe frequency within the bottom PDCUB quintile. Triangles by the labels indicate enriched (pointed up) and depleted (down) GO terms. **B.** Scatterplot representing statistically enriched GO terms associated with transcripts in the highest PDCUB score quintiles, displayed similarly as A. **C.** Local tAI profiles for transcripts in quintiles defined for PDCUB score. **D.** The shape of ribosomal occupancy, shown as the normalized Ribo-seq profile centered at the start codon at position 0 for quintiles defined for PDCUB. Read positions are A-shifted by 15-nt, and smoothed using a Savitzky-Golay filter.

##### Growth and development

Among the highest-PDCUB transcripts, we observed a statistical enrichment of most developmental processes, with a corresponding depletion of these same processes among the lowest-PDCUB transcripts (Supplementary Table 1, blue terms). Examples include generation, development, differentiation, formation, and morphogenesis of the sensory, nervous, endocrine, gastrointestinal, circulatory, and musculoskeletal systems, as well as of various organs, glands, tissues, cells, and cellular components. Terms with the greatest enrichment (> 5*x* expected) were nephron tubule formation and regulation of norepinephrine secretion, the latter of which is increasingly thought to be a uniquely important regulator of brain development [17]. The only exceptions to this trend were development-related terms involving skin and keratin, which were enriched only among low-PDCUB transcripts.

##### Signaling and transport

Nearly all significant GO terms pertaining to signaling and transport are enriched among high-PDCUB transcripts but depleted among mid- and low-PDCUB transcripts (Supplementary Table 1, purple terms). Exceptions to this trend primarily consist of terms tied to immunological function, which are depleted among high-PDCUB transcripts but enriched among mid- and low-PDCUB transcripts, as well as G-protein coupled receptor signaling pathway terms that are enriched across all PDCUB categories.

##### Immune response and defense

Significant GO terms pertaining to immune response are depleted among high-PDCUB transcripts and enriched among mid- and low-PDCUB transcripts (Supplementary Table 1, green terms). We found that “adaptive immune response” was the most statistically significant GO term in the analysis of both the significant PDCUB set and the top PDCUB quintile, and was strongly depleted in both cases. We found a single reversal of this trend in the term “phagocytosis”, which is the most depleted term in the second PDCUB quintile. Interestingly, terms relating to specific immune cells such as lymphocytes, leukocytes, granulocytes, monocytes, eosinophils, neutrophils, and natural killer cells are highly enriched among the middle quintile (40%-60% PDCUB scores) but neither significantly enriched nor depleted in the other quintiles.

##### Chemical stimulus and sensory perception

Significant GO terms related to stimulus *response* and *regulation* are all *enriched* among high-PDCUB transcripts (Supplementary Table 1, pink terms). By contrast, significant GO terms pertaining to sensory or stimulus *detection* and *perception* are all *depleted* among the high-PDCUB transcripts (Supplementary Table 1, yellow terms).

Among mid- and low-PDCUB transcripts the opposite situation holds true, with enrichment of GO terms involving sensory or stimulus detection and perception, and depletion of terms involving stimulus response and regulation.

#### Signal Peptides

To investigate whether our results reflect the presence of nucleotide sequences coding for signal peptides, we used the signal-peptide prediction tools PrediSi [18] and SignalP [19] to predict which transcripts are likely to contain signal peptides. We performed a statistical test comparing the number of predicted signal peptides in the high-significance PDCUB transcripts to a background predicted rate for a complement set of all protein-coding GENCODE transcripts not contained within the significant PDCUB set. We observe that 15-24% of proteins encoded by high-PDCUB transcripts are predicted to contain signal peptides, amounting to a 1-4% increase in predicted signal peptides among high-significance PDCUB transcripts relative to all proteincoding transcripts in GENCODE (Supplementary Figure 5). We tested this difference via binomial test and found the increase to be statistically significant, with a p-value of < 10^−23^. A Venn diagram of transcripts predicted by each approach to have a signal peptide, compared to transcripts with a high-significance PDCUB score, is shown in Supplementary Figure 6. Analysis of individual PDCUB quintiles showed a rough correlation between signal peptide density and quintile number for PrediSi and SignalP alike. In both cases, we found the bottom quintile to have the lowest predicted density, with the third quintile being near the global average density and the fourth quintile having the highest predicted density (Supplementary Figure 7).

### Local tAI and CAI

We next sought to contextualize PDCUB against standard measures of translational efficiency, including the discovery of the translation initiation ramp [20]. We confirmed the presence of the translational ramp seen by Tuller et al, through a metagene plot of tAI against CDS position for the first 300 codons of the coding transcriptome (Supplementary Figure 8). We repeated this process with the transcriptome split into quintiles according to PDCUB scores, and found a clear, nonoverlapping spectrum of ascending average tAI values as we move from the lowest-PDCUB transcripts to the highest-PDCUB transcripts (Figure 4C). This PDCUB-defined spectrum exhibits differently shaped local tAI curves, with the fifth quintile (top quintile, highest PDCUB score) having the most pronounced translational ramp and lower quintiles having diminishing ramp size until the second quintile, which is flat with no ramp. More strikingly, the first quintile (bottom quintile, lowest PDCUB score) exhibits a “translational valley”, with tAI descending quickly from codons 0 to 30 before flattening out until codon 100, whereupon we observe a short ramp that is delayed relative to that of the highest PDCUB quintile.

We also compared tAI trends for quintiles defined for CAI computed for the entire coding sequence (global CAI), as well as for specific regions of the coding sequence (regional CAI). Trends for tAI plotted against quintiles defined for global CAI and regional CAI for the first 100 codons showed similar but much weaker trends, with global CAI showing the weakest changes in tAI between the three methods (Supplementary Figure 9). Quantitative comparison of quintile-specific tAI trajectories between the three methods confirmed that for both PDCUB and regional CAI the greatest change in tAI was seen in the first and fifth quintiles, corresponding respectively to a translational valley and a strong ramp, while global CAI quintiles display a steady monotonic increase from quintiles one through five (Supplementary Figure 10). Finally, we plotted tAI against PDCUB score for the coding transcriptome and found poor general correlation between the two parameters (*R*^2^ ≈ 0.3, Supplementary Figure 11).

We calculated local tAI values for CDS regions corresponding to the first and second windows of length *w* codons, for *w* = 30 and *w* = 50, for all transcripts. The midpoints of these two sets of paired windows correspond, respectively, to the lower and upper bounds reported for the size of the translation initiation ramp and should thus demark a boundary between lower and higher translational efficiency signals, which should be apparent when plotting the local tAIs for the second window against the first. No such boundary was apparent in our scatter plot, regardless of the PDCUB of the corresponding transcripts (Supplementary Figure 12). However, we did see a clear gradient of PDCUB scores across the transcript distribution in both codon windows, with low-PDCUB transcripts concentrated at low tAI values, and high-PDCUB transcripts concentrated at high tAI values.

As an alternative approach to understanding how PDCUB compares to translational efficiency, we performed a similar set of analyses with regional CAI scores computed from global and regional relative adaptiveness (Supplementary Figures 13 and 14). This revealed a trend in which transcripts with lower PDCUB scores correspond to a diminished regional CAI in codons 30-50 (the translation initiation ramp) relative to the immediately subsequent 30-50 codons (the translation steady state). We plotted the CAI of each protein-coding transcript against its PDCUB score, and found the relationship between CAI and PDCUB to be similar to (but weaker than) that between tAI and PDCUB (Supplementary Figure 15).

We further compared the CAI for the first 100 codons to score AUGs for the prediction of start codons in the same analysis as Figure 3A. We found that CAI was not predictive for start codons, and was in fact lower than random for the same window that PDCUB is computed (Supplementary Figure 16).

### Ribo-seq profiles

Given the observation that higher ribosomal density is associated with lower translational speed [21], we examined whether transcripts with higher PDCUB scores show distinct trends in ribosomal occupancy compared to transcripts with lower PDCUB scores. Figure 4D shows the normalized ribosomal occupancy relative to the start codon for each PDCUB quintile. We observed a spectrum of ribosome profiles where the first quintile (lowest PDCUB) has the highest relative occupancy at initiation while the fifth quintile (highest PDCUB) has the lowest relative occupancy. In contrast, a control experiment of ribosomal density plotted against CAI quintiles showed no such clear trend between the two metrics (Supplementary Figure 17).

## DISCUSSION

We observe position-dependent trends when visualizing codon usage across all protein-coding regions in the human transcriptome. Additionally, over one third of ORFs in individual proteincoding transcripts exhibit a statistically significant PDCUB when compared to reading frames that follow non-start AUGs. We have shown that start codons can be distinguished from other AUGs across reading frames solely by using a score that quantifies PDCUB for the 300 nt immediately following the AUG, thus further validating that PDCUB is a distinguishing feature of many start codons. We also observe a substantial difference between the ribosomal occupancy for higher-PDCUB transcripts compared to those with lower PDCUB. These observations are consistent with the hypothesis that human translational machinery, including ribosomes and translation initiation factors, rely broadly upon positionally dependent sequence patterns to direct and regulate multiple aspects of translation.

As discussed in previous studies in yeast [13], we tested whether our observed PDCUB pattern could be explained by signal peptides and found a 1-4% enrichment of signal peptides in proteins encoded by high-significance PDCUB transcripts (Supplementary Figure 5). Sorting predictions by PDCUB quintile revealed a rough correlation between PDCUB quintile and predicted signal peptide density, but even the most overrepresented quintile showed < 5% enrichment in signal peptides relative to the global background (Supplementary Figure 7). Interestingly, despite this correlation, both SignalP and PrediSi predict a maximum overrepresentation of signal peptides in the fourth rather than fifth PDCUB quintile. This might be explained by the fourth quintile being uniquely composed of a single overrepresented GO term related to GPCRs, which are known to have a high (≥ 5%) rate of occurrence of signal peptides [22]. Overall, the low predicted enrichment of signal peptides in all PDCUB quintiles suggests that while sequence features coding for signal peptides may contribute slightly to PDCUB, they are not its driving mechanism. It is also worth noting that while signal peptides are defined by an amino acid sequence, PDCUB describes position-dependent patterns at the nucleotide level in addition to the triplet level, and thus may capture translationally important patterns that position-dependent amino acid signals alone cannot completely resolve.

Previous research has shown that GC-rich codons have greater translational efficiency [23]. We observe that the pattern of PDCUB across transcripts shows enrichment of GC content in codons at the beginning of coding regions relative to downstream segments. Simultaneously, we observe lower ribosomal occupancy over the region in which PDCUB is calculated (1-300 nt downstream of the start codon). Therefore, we speculate that observed PDCUB patterns may lead to greater efficiency of translation initiation and possibly also translation elongation. Selection for GC-rich, translationally efficient coding regions may encourage the development of PDCUB patterns, or vice versa. However, the fact that PDCUB outperforms GC content in the identification of start codons suggests that GC content is not the whole story, and higher-order sequence patterns may be at play.

We observed that PDCUB defines a spectrum of translational efficiency, as seen in both ribosome occupancy and local tAI trajectories. Previous studies have been conflicted about the presence of a translational ramp in mammals, with some reporting no ramp [24] and others reporting a distinct ramp [20]. We find that both results are true depending on where one looks within the PDCUB spectrum, with the highest-PDCUB transcripts having a strong initial ramp, intermediate PDCUB transcripts having a weaker ramp or no ramp, and the lowest-PDCUB transcripts exhibiting an initial translational valley that, to our knowledge, has not been described before. In particular, high-PDCUB transcripts seem to exhibit the same translational ramp described elsewhere [20, 25], and this translational ramp corresponds to reduced ribosomal occupancy in the early regions of those same transcripts. In contrast, tAI trajectories plotted against global CAI quintiles show a much weaker ramp and no valley (Supplementary Figure 9). When we alter the CAI calculation to only include the first 100 codons of each transcript (the same window in which PDCUB is calculated) the altered trajectories display a much weaker valley (more of a delayed ramp) and an initial ramp profile that begins to more closely (but still weakly) resemble PDCUB, while downstream regions display poor separation compared to their PDCUB and global CAI equivalents. All of this indicates that PDCUB dramatically improves on CAI in its ability to detect important early regions of variable translation efficiency, while retaining the ability of CAI to clearly identify stable downstream regimes of translational efficiency – a balance which modified regional CAI fails to achieve, leaving PDCUB as the best of the three options. The low start codon predictive potential of CAI computed from the first 100 codons is further evidence of the limited ability of CAI to capture the dynamic patterns of codon usage at the start of coding regions.

Upon quantifying the absolute growth of these tAI trajectories for PDCUB, CAI, and regional CAI quintiles, we see that distinct trends occur for each measure of coding potential (Supplementary Figure 10). Transcripts in the highest PDCUB quintile show the greatest growth in localized tAI, while the second-greatest growth occurs for the lowest quintile, corresponding to emergence from the translational valley. By contrast, for standard CAI we see a monotonic increase in the height of the localized tAI growth across all quintiles, due to a lack of a translational valley for the lowest CAI quintile. This is further evidence that PDCUB captures important information regarding translational efficiency that is not captured by standard applications of CAI and tAI alone, as reinforced by the low direct correlation between these metrics over the entire protein-coding transcriptome (Supplementary Figures 11-15).

Importantly, the PDCUB-defined transcript subpopulations correspond to transcript function (Supplementary Table 1, PDCUB sheets). In our analysis of GO terms, we observe that functions involved in growth and development at the system, organ, tissue, cellular, and subcellular levels are highly enriched among high-PDCUB transcripts. Among low-PDCUB transcripts, such terms are depleted. This could be explained by the fact that developmental pathways are part of precise regulatory functions that require exquisite tuning of translational control [26, 27] and precise timing [28, 29], and that PDCUB may serve as a mechanism to enhance translational efficiency within regulatory cascades. This may be true particularly at the translation initiation step, thereby accounting for the pronounced translational ramp found among high-PDCUB transcripts and the enhanced ability of PDCUB to detect start codons.

The only exceptions to the trends described above are for developmental terms involving skin and keratin, which are enriched among low-PDCUB transcripts only. This set of exceptions may be tied to the prominent role of these tissues in human immune function relative to other organs and tissues, as they comprise the body’s first and primary barrier against pathogens. More generally, we observe that transcripts related to immune response, as well as those governing detection and perception of chemical stimuli, are statistically depleted among high-PDCUB transcripts and enriched among low-PDCUB transcripts. Immune response and chemical detection pathways have in common high genetic copy number and sequence variation [30]. Moreover, there is an abundance of pseudogenes observed for human odorant receptors and other genes responsible for *detection* and *perception* of chemical stimuli [31], as well as immune genes [32, 33]. These observations may suggest a lower selective pressure on individual proteincoding genes due to the robustness of these pathways conferred by high copy number and a need to constantly adapt to changing chemical [34] and pathogen [35–37] environments. We posit that genes within these pathways might sample a broader range of codon ordering compared to genes involved in the development of complex vital organs. Meanwhile, genes tied to chemical stimulus *response* and *regulation* show a reversed trend relative to detection and perception, with enrichment among high-PDCUB transcripts and depletion among low-PDCUB transcripts. This trend reversal may stem from a need for precise timing among regulatory and response genes, as described for developmental genes above, while detection and perception genes rely on high copy number and low translational control to account for the complex suite of chemicals that must be adapted to in the human sensory environment.

Comparison of these trends to those for CAI quintiles reveals at best partial qualitative similarity for some of the overarching GO categories identified above, with several interesting and significant deviations (Supplementary Table 1, CAI sheets). Most notably, we see enrichment of immune terms among high-CAI transcripts and depletion among mid- and low-CAI transcripts – exactly opposite of the trend seen for PDCUB quintiles. A similar trend reversal is seen for terms relating to skin, which are enriched exclusively in the highest CAI quintile and the lowest PDCUB quintile. Given the prominence of these trend reversals, and paired with the critical role of immune response in human evolution, we postulate that PDCUB and CAI play functionally distinct roles in translational regulation. For instance, it is possible that CAI governs steady-state baseline levels of background translation, with high CAI reserved for those genes that are more consistently active, while high PDCUB correlates to situationally sensitive genes that require precise timing and a more well-defined translational ramp. While these two categories might overlap (as in the development terms), it is also possible for them to diverge sharply (as in the immune terms). Additional evidence for this view is the presence of biologically cyclic GO terms such as “hair cycle” and “molting cycle” in the results of the CAI quintile analysis, but the total absence of such terms from the PDCUB analysis. Further evidence lies in the split described above for stimulus and sensory GO terms. Among PDCUB quintiles the relationship between these terms is neatly split between low- and high-PDCUB transcripts, with detection and perception showing enrichment among low-PDCUB transcripts and depletion among high-PDCUB transcripts, while regulation and response follow the opposite trend. In contrast, CAI quintiles do not show this split and instead have detection and perception terms uniformly enriched, but response and regulation terms uniformly depleted, across all quintiles in which they appear (the exception being quintile 4, which has no stimulus or sensory terms). These results serve as a basis for understanding the complex relationship between PDCUB and other metrics of codon bias, including the extent to which they might be complementary or competing factors in translational efficiency.

## MATERIALS AND METHODS

The dataset used was the GENCODE Release 34 protein-coding transcript sequences FASTA [15]. We filtered out all transcripts in which the length of the labelled coding region was not a multiple of three, lacked an in-frame start codon, or lacked an in-frame stop codon. This dataset was used for all of our analyses.

### Codon frequencies relative to transcript sequence position

Codons following the start codon were counted in bins of size 60 (20 nucleotides) for a total of 50 bins. To keep the bin size constant, the probability of finding a given codon in a bin was calculated, and subsequently used to derive a mean and standard deviation for each codon. The z-score of a given sense codon in a given bin in Figure 1A was calculated by

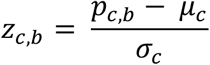

where *p_cb_* is the probability of a codon in a given bin, *μ_c_* is the mean of the codon across all bins, and *σ_c_* is the standard deviation of the codon across all bins.

### Hierarchical clustering of codons based on z-score profiles

To obtain the dendrogram of z-score profiles, a distance matrix was first calculated wherein each row and column corresponds to a single codon. A cell at row *m* and column *n* would thus correspond to the distance between the z-score profiles of codons *m* and *n*. The distance between two codons was calculated based on the sum of squares of the differences between z-scores:

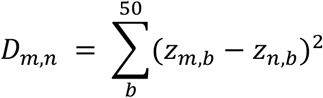

where *Z_mb_* and *Z_nb_* are respectively the z-scores of codons *m* and *n* at bin b.

The dendrogram was computed using python’s scipy.cluster.hierarchy.linkage method [38], which performs clustering given a distance matrix such as the one calculated above. The method used for calculating distance between two clusters *u* and *v* was the ‘average’ method

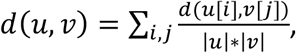

which describes the distance between two clusters of codon profiles over positions *i* and *j* relative to the cardinalities of clusters *u* and *v*.

### Scoring models for predicting start codons

Several scoring models were developed to predict the location of start codons, in order to determine the effect of position dependence in the human transcriptome. These models were based on a weight matrix, or position specific-scoring matrix (PSSM), where the rows correspond to codons or nucleotides and the columns correspond to positions after the start codon. In general, the PSSM values were calculated as log likelihoods based on a background model.

#### PDCUB PSSM

The weights in the PDCUB PSSM were calculating according to the equation

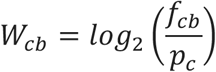

where *f_cb_* is the frequency of codon *c* at bin *b*, and *p_c_* refers to the background probability of codon *c*. Bins *b* of size *w* (in codons) with start positions *s_b_* comprise a series {[*s*_1_, *s*_1_, + *w*], [*s*_2_, *s*_2_ + *w*],… [*s_N_, s_N_* + *w*]} up to *N* bins for each putative CDS. We define a putative CDS region as series of triplets after the AUG, *C* = (*c*_1_, *c*_2_,…, *c_n_*). The PDCUB score *S*(*C*) is computed for a given CDS according to

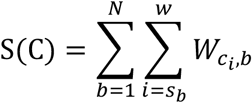

with PDCUB models computed at bin sizes of *w* = 1, 2, 3, 4, 5, 10, 20, 25, 50, and 100 codons. After examining the percentage of transcripts in which the true start codon was the highest scoring AUG, it was determined that a bin size of 3 nucleotides was optimal.

#### Kozak consensus sequence

The Kozak consensus sequence is defined as (gcc)gccRccAUGG, where R corresponds to A or G, and position +1 corresponds to the A in the AUG. Positions −3 (the R) and +3 (the G after the AUG) are particularly important, and upper-case indicates highly conserved bases [16]. We gave positions +3 and −3 a weight of 6, and the remaining positions a weight of 1. The score of +6 for each highly conserved position ensures that correctly matching one highly conserved position outweighs any number of less-conserved matches. Transcripts were scored based on the number of mismatches the surrounding nucleotides had with the consensus sequence, with a maximum score being 17 (no mismatches). The leading (gcc) was subtracted from this score, so that codons containing the matching sequences were rewarded, but codons without them were not penalized.

#### Kozak PSSM

The Kozak PSSM model was implemented in a manner similar to that for PDCUB, with a few key differences. A PSSM was built by computing the frequencies *f_b,i_* for each nucleotide *b* at position *i* relative to the start codon across all protein-coding transcripts. The PSSM is then computed using this frequency relative to the global single-nucleotide frequencies *p_b_*, as shown below.

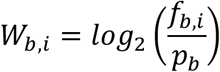

Therefore, the Kozak PSSM differs from the PDCUB PSSM because the former was based on nucleotide counts instead of codon counts. The positions in the Kozak PSSM correspond to the ten nucleotides before the start codon, which is the location of the Kozak consensus sequence. All positions in the transcript were thus scored by taking the sum of the weights of ten preceding nucleotides.

#### GC content PSSM

The GC content PSSM has two rows, one corresponding to ‘S’ nucleotides {G,C} and the other to ‘W’ nucleotides {A,U}. Nucleotides were counted into either of those categories in a manner similar to the nucleotide PSSM used for the Kozak weight matrix.

#### Statistical Null Model

The null model was calculated to determine the apparent extent of random chance involved in an AUG being the true start codon in a given transcript. All AUG triplets in a transcript were scored, and an AUG was randomly selected as the start codon. If the selected AUG had the highest score, then the model was considered successful for that transcript.

#### Theoretical Model

If the length of a given transcript is *L*, then there are *L* – 2 possible positions in which a triplet can exist when allowing all three possible reading frames. Moreover, the codon at that given position must be an AUG, of which there is a 1/64 chance. Therefore, the total number of expected AUGs in a transcript is

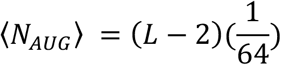

The theoretical probability that an AUG selected a random is the true start codon can thus be modeled by

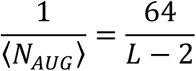

### Cross-validation methods

The dataset of all valid sequences was split into three separate sets. In all cases, 80% of the data (about 55,000 transcripts) were used to obtain PSSMs of the model in question. In most cases, the remaining 20% (about 14,000 transcripts) were scored with the calculated PSSM. In the case of PDCUB, where multiple bin sizes were tested, 10% of the data was used to determine the bin size that produced the highest-scoring AUG triplets, and 10% was left for scoring.

### Information content of bins within a transcript

The information content *IC* of position *i* in a transcript was calculated as follows:

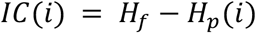

where *H_f_* is the Shannon entropy of the global triplet frequencies (background model) of the transcript, and *H_p_*(*i*) is the Shannon entropy of the position-dependent codon frequencies at position *i*. The background entropy is calculated as

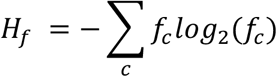

where *f_c_* is the global frequency of triplet *c*. The product of the global frequency and the log of the global frequency are computed for all triplets and then added together. The Shannon entropy of the position-dependent codon frequencies is calculated as

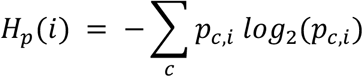

where *p_ci_* is the probability of codon *c* at position *i*.

### Determination of high-significance AUG triplets and PDCUB quintiles

Upon generating a list of transcripts and their highest-scoring AUG triplet based on the PDCUB model, the p-values for each AUG score were calculated. All scores were sorted by p-value and accordingly ranked (the lowest p-value would have a rank of 1, the next lowest a rank of 2, and so on). These values were filtered with a Benjamin-Hochberg multiple test correction to produce a list of statistically significant genes (*α* < 0.05), with corresponding q-values calculated according to

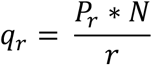

where *r* is the rank and *N* is the total number of scores. All transcripts with a rank lower than the highest rank *r* with *q_r_* below the significance level were deemed statistically significant. This set was considered the high-significance PDCUB set (significant PDCUB set), while an equivalently sized transcript set with the highest *q_r_* was considered the low-significance PDCUB set.

A second ranking system was also used, wherein all transcripts were divided into a set of quintiles according to PDCUB score. The 20% of transcripts with the highest PDCUB score comprise the top PDCUB quintile and so on, in Table 1.

**Table 1:** Definition of PDCUB quintiles.

| PDCUB quintile | percentile range (%) | quintile name |
| --- | --- | --- |
| 5 | 81-100 | fifth quintile, top quintile, highest PDCUB |
| 4 | 61-80 | fourth quintile |
| 3 | 41-60 | third quintile, middle quintile, moderate PDCUB |
| 2 | 21-40 | second quintile |
| 1 | 01-20 | first quintile, bottom quintile, lowest PDCUB |

### Analyzing the contribution of signal peptides

#### Prediction of signal peptides in human proteome

The mRNA content of the human transcriptome was retrieved from GENCODE, converted to protein sequence format, and fed to the online tools PrediSi [18] and SignalP-5.0 [19] for prediction of signal peptides. Both tools were accessed through their public-facing online interfaces using default program parameters. In both cases, transcripts from the significant PDCUB set were separated from the remainder of the GENCODE set.

#### Statistical enrichment of signal peptides in significant PDCUB set and quintiles

Predicted signal peptide densities for the significant PDCUB set and each PDCUB quintile were compared to background signal peptide enrichment in two batches: one for the PrediSi scores, and another for the SignalP scores. In both cases, the significant PDCUB set and quintile sets were tested for enrichment of signal peptides relative to the global set. Enrichment significance was determined via binomial test.

#### Overlap between significant PDCUB and signal peptides

Transcripts from the significant PDCUB set were checked for overlap with all GENCODE proteincoding transcripts predicted to have a signal peptide. This was done for two sets of transcripts: GENCODE transcripts that scored positively for signal peptides using PrediSi, and those that scored positively for signal peptides using SignalP-5.0. All three resulting sets (significant PDCUB, PrediSi GENCODE, and SignalP-5.0 GENCODE) were visualized together as a Venn diagram using matplotlib.

### Gene Ontology Enrichment

ENSEMBL IDs for transcripts were analyzed for each PDCUB category (high-significance, top quintile, and so on) using the GO Enrichment Analysis tool, which is part of the web-accessible Gene Ontology Resource (http://geneontology.org/) [39–41]. Tables of statistically significant GO terms were retrieved for both sets of transcripts. The tables are provided as excel files in Supplementary Table 1. Significant GO terms were those with unexpected enrichment or depletion among a given PDCUB category, when a Benjamini-Hochberg multiple test correction with a false discovery rate of 0.05 was applied.

### Calculating regional CAI and local tAI

Sets of local relative adaptiveness (RA) and CAI values were calculated for all protein-coding GENCODE human transcripts. Whereas RA weights are traditionally defined by one unique set of values for a given transcriptome, and are derived from the total codon composition across the full lengths of every CDS, our local RA weights were computed for a particular region r of every CDS. These local RA values were used as input weights *w_rc_* to calculate regional CAI values for the same region r and codon *c* of a single transcript. For example, when examining the regional CAI of the first 50 codons in a given CDS, our local RA was derived exclusively from the first 50 codons of every CDS. The local RA was calculated as

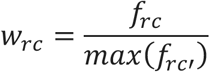

where *f_rc_* is the frequency of codon *c* in region *r* and *f_rc′_* is the frequency of codon *c′* in region r, where *c* and *c′* are both synonymous codons for the same amino acid. From this, the regional CAI was calculated

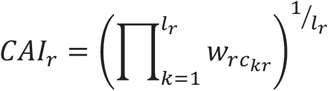

as where *l_r_* is the length of the transcript region, and *c_kr_* is the codon defined by the *k*^th^ triplet in that region.

To calculate local tAI, we retrieved human codon tAI weights from Table S1 of Tuller *et al*. [20] and computed a similar geometric mean for the same regions as above. Quintile tAI figures were generated by splitting GENCODE transcripts into five parts sorted by PDCUB score. Unlike regional CAI, which consists of a single average value for the entire region under consideration, average local tAI was plotted for each quintile as a function of codon position using a sliding window five codons in length.

### Ribosome profiling

Ribosome profiling of HEK293 cells was downloaded from GSE accession number GSE102720 [42]. Ribo-seq reads were aligned to protein-coding transcripts using Bowtie [43] with the parameters “-v 2 -a -S --strata --best --norc -m 200” after trimming the adaptors with Cutadapt [44]. Read 5’ ends were shifted relative to the start of the CDS and an additional 15 nucleotides to align to the A site [45]. Normalization of each individual transcript was performed by dividing the sum of the reads at each nucleotide position by the total number of reads mapping up to 500 codons from the start codon for that transcript, and converted to a percentage. The sum at each position of these normalized transcripts was divided by the total number of transcripts to create an average profile. The average curves for each quintile were smoothed using a Savitzky-Golay filter [46] with a window size of 19 and a polynomial degree of 4 using the scipy implementation [38].

### Project code

Code for this project is available at https://github.com/hendrixlab/PDCUB.

