## Supplementary Materials for "Position-dependent Codon Usage Bias in the Human Transcriptome"

1. School of Electrical Engineering and Computer Science, Oregon State University
2. Department of Biochemistry and Biophysics, Oregon State University
* these authors contributed equally to this work

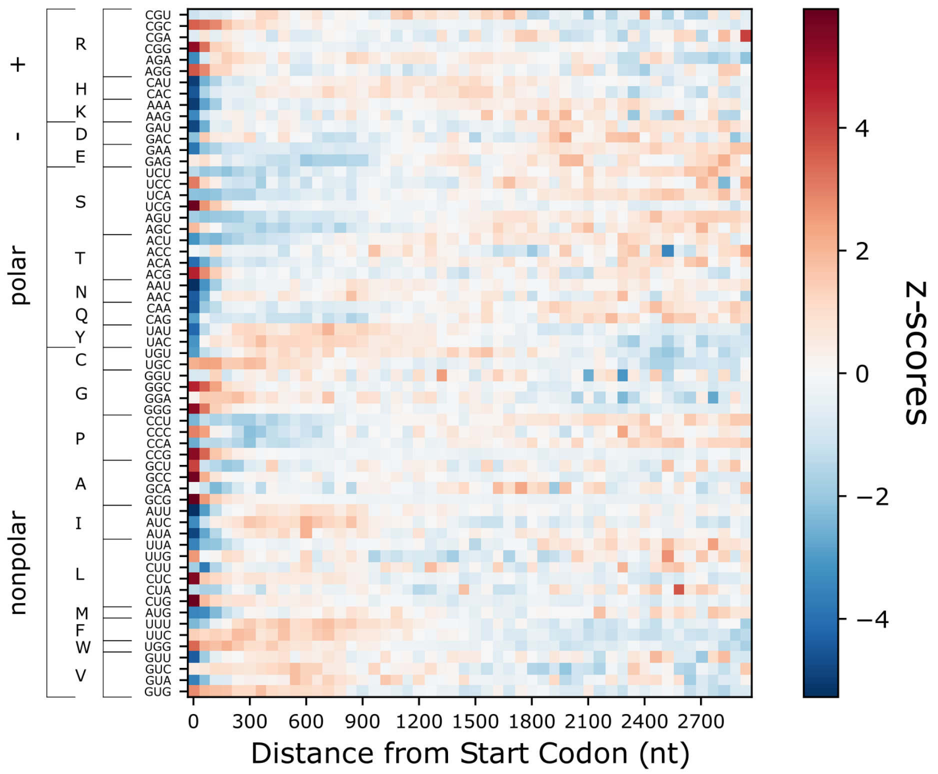


***Supplementary Figure 1:*** *Z-score heatmap computed from codon counts in bins of size 60 nt, for the first 3000 nt of all coding regions. Codons are grouped by amino acid and by the chemical category of the amino acid.*

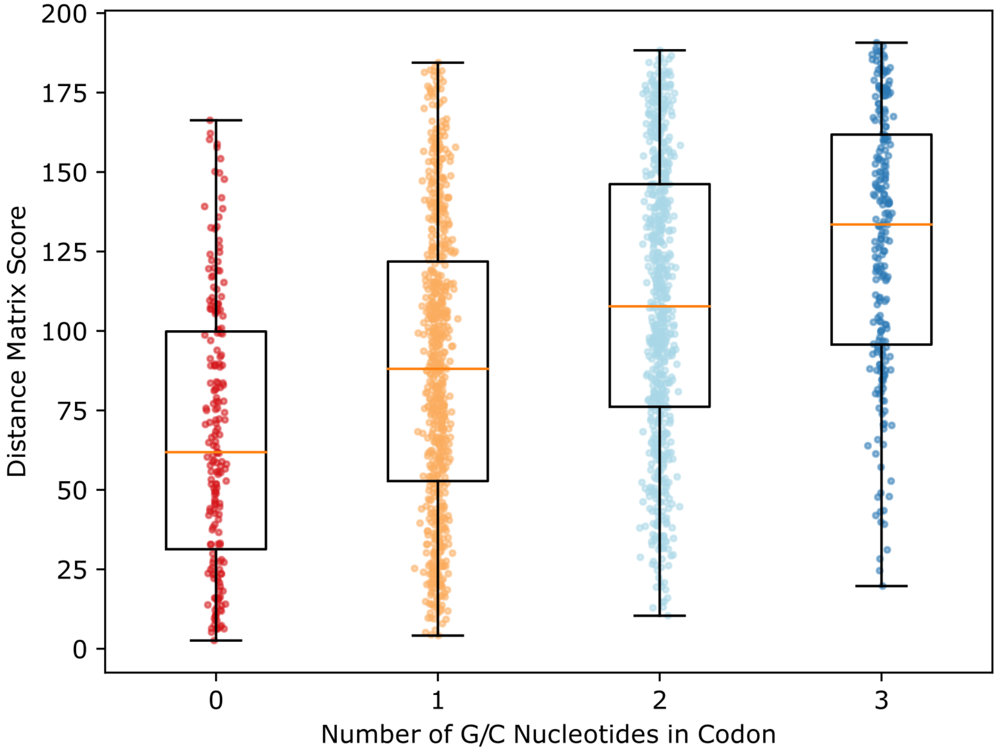


***Supplementary Figure 2:*** *Scatter-boxplot comparing the distance between codons based on the difference in GC nucleotides versus distance between codons based on hierarchical clustering of the position-dependent z-score profiles.*

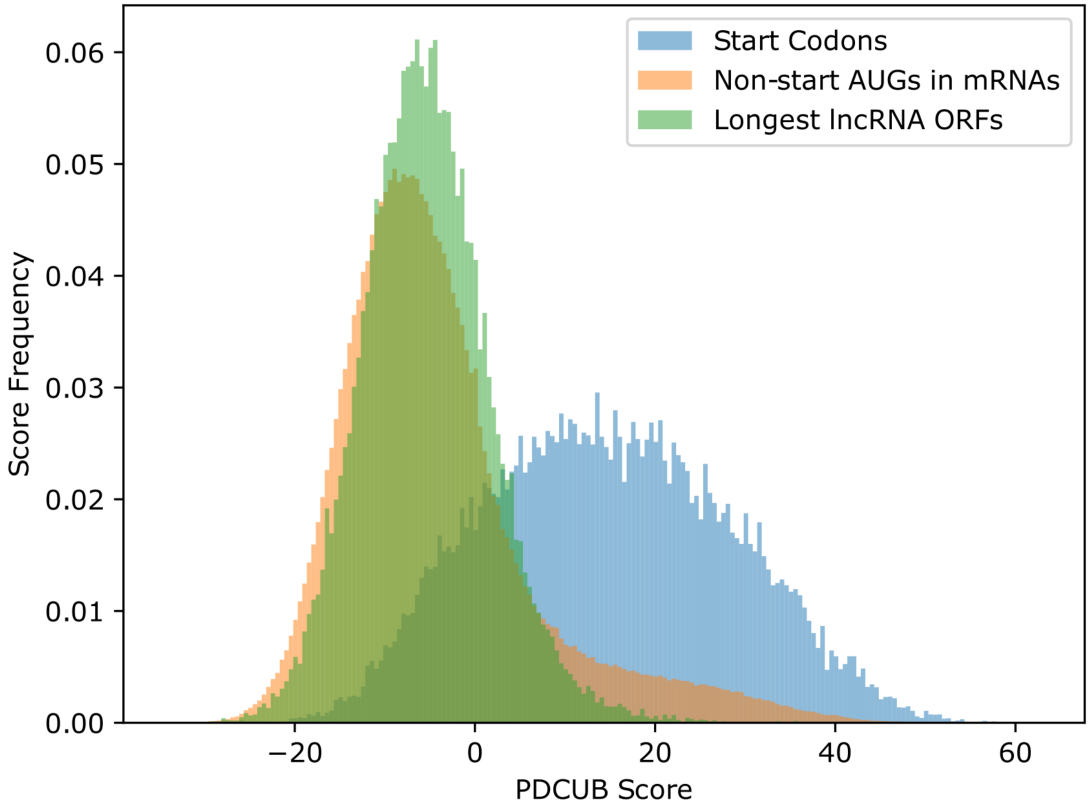


***<u>Supplementary Figure 3:</u>*** *Histograms comparing the PDCUB score for start codons compared to non-start AUG triplets and the AUG in the longest ORF for lncRNAs. Mean and standard deviation from the non-start distribution is used to compute statistical significance of PDCUB scores.*

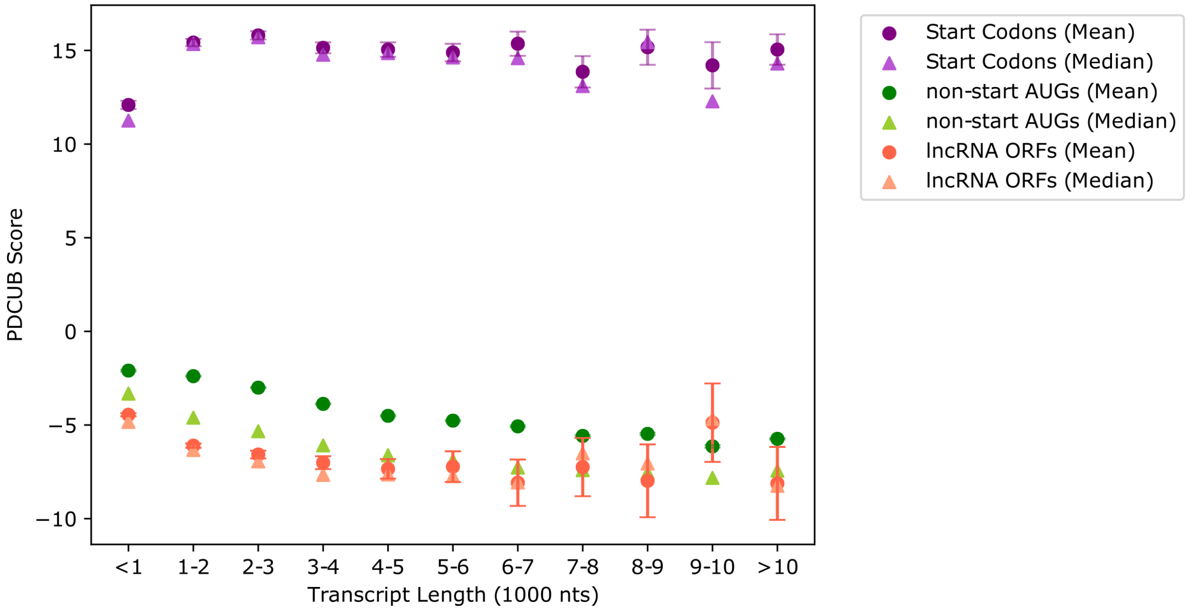


***Supplementary Figure 4:*** *Comparison of the mean and median PDCUB score for start codons in transcripts of different lengths in the test set, relative to non-start AUG triplets, in increments of 1000 nt. In addition, this also compares to the scores for the AUG in the longest ORF for lncRNAs using the same size bins.*

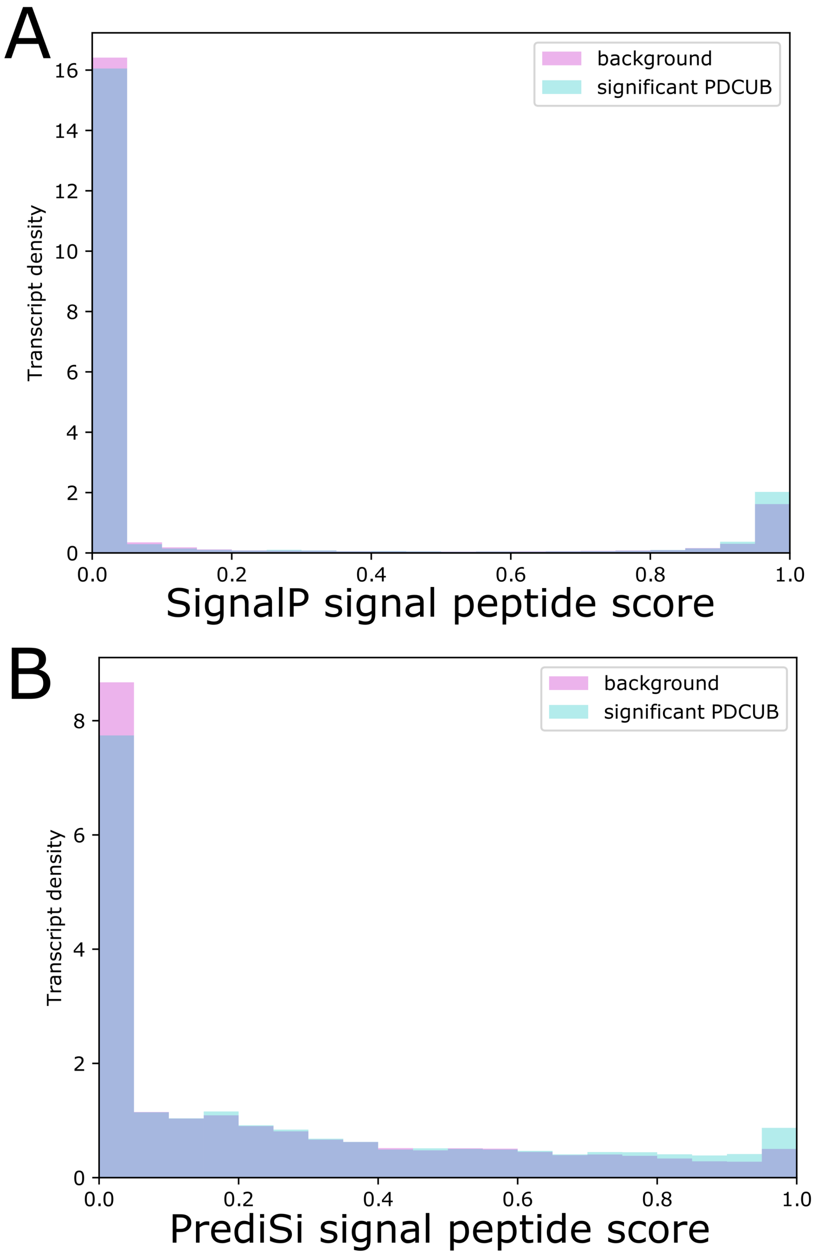


***<u>Supplementary Figure 5:</u>*** *Histograms comparing the predicted probabilities of signal peptides for significant PDCUB transcripts against those of the full GENCODE set.* ***A.*** *SignalP distribution.* ***B.*** *PrediSi distribution.*

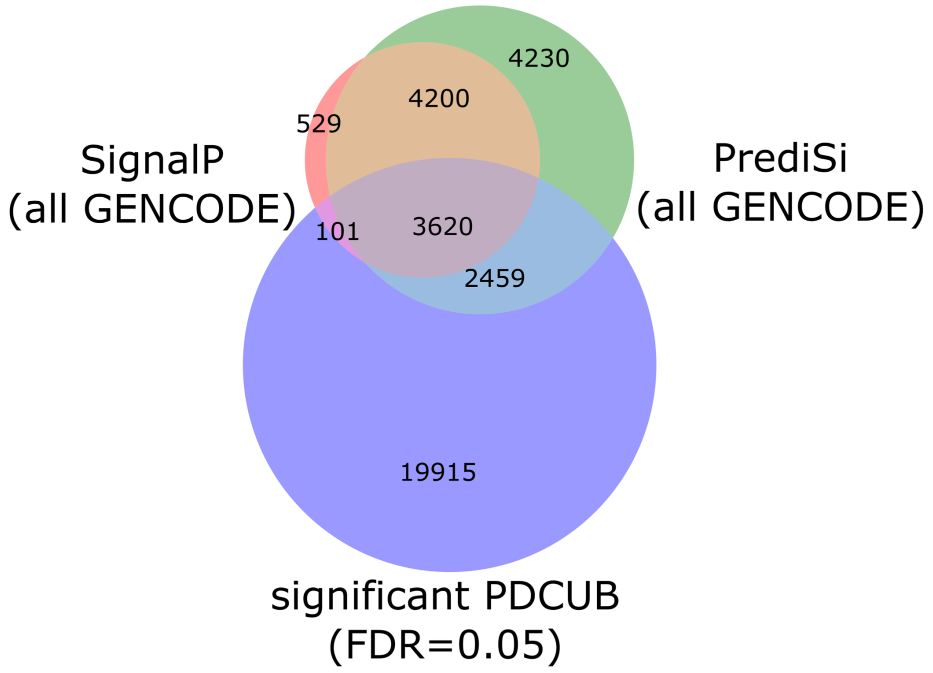


***Supplementary Figure 6:*** *Venn diagram of coding transcripts predicted to have signal peptides according to SignalP (top left) and PrediSi (top right), compared to coding transcripts that meet the significance threshold for high PDCUB score (bottom).*

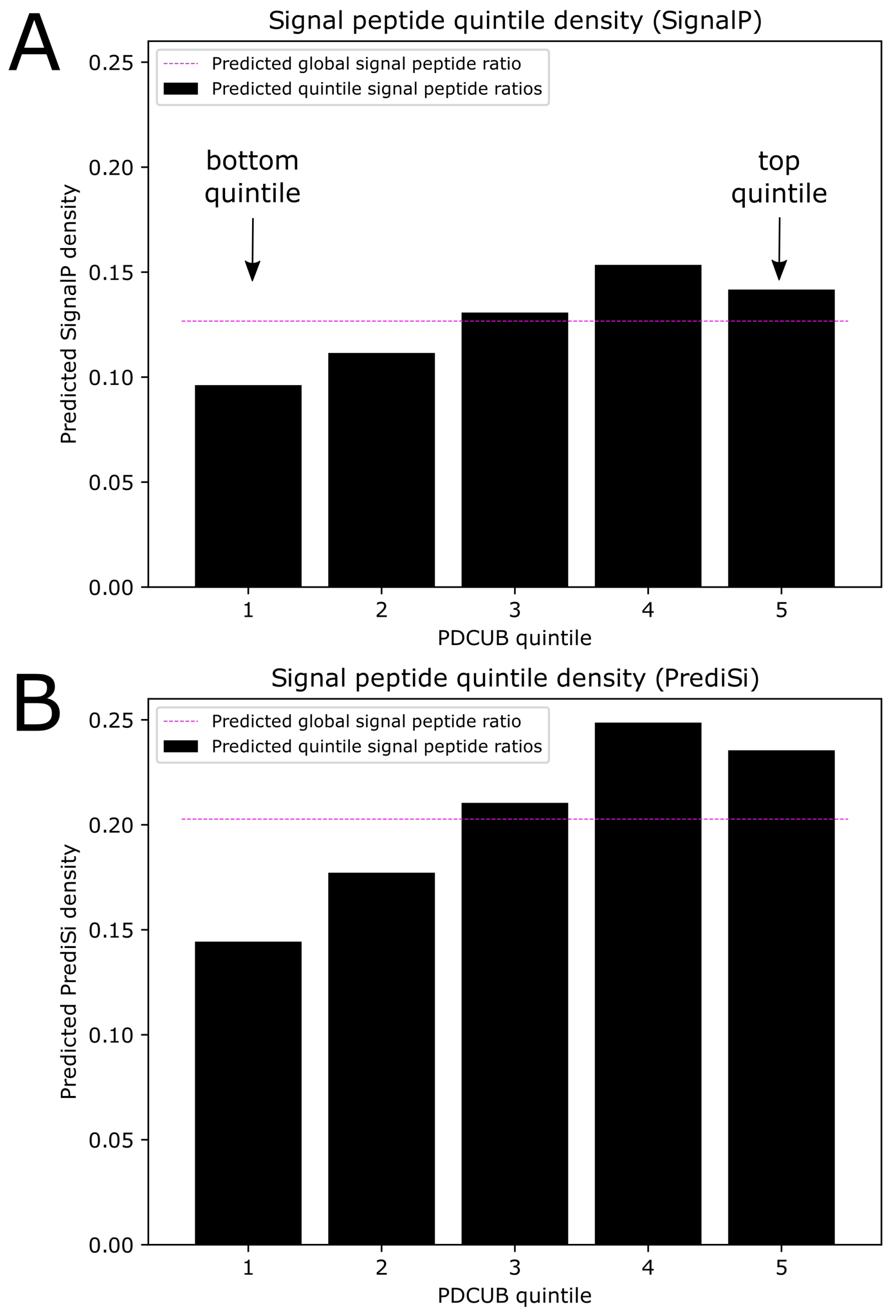


***Supplementary Figure 7:*** *Predicted signal peptide densities for each PDCUB quintile. Magenta line indicates global predicted signal peptide density.* ***A.*** *SignalP predictions.* ***B.*** *PrediSi predictions.*

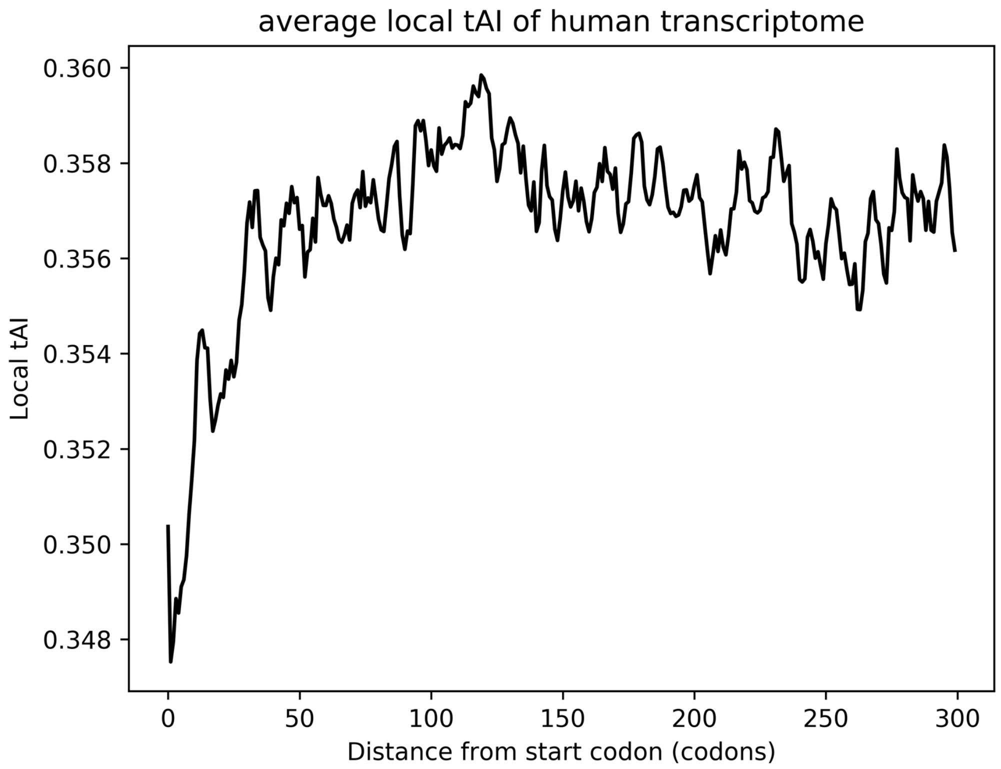


***Supplementary Figure 8:*** *Average local tAI for first 300 codons of coding transcriptome in humans, using a sliding window of size 5 codons. A distinct ramp is seen from codons 0 to 50.*

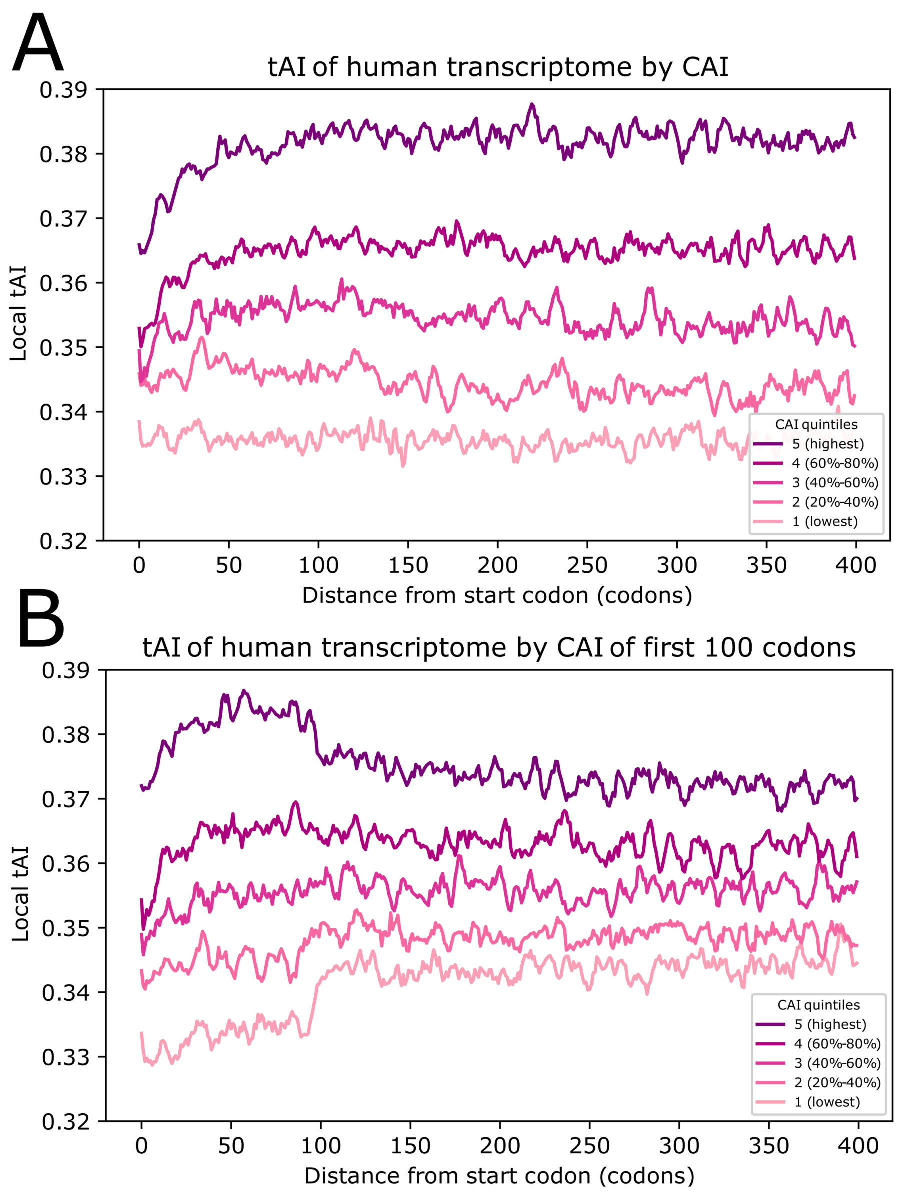


***Supplementary Figure 9:*** *Local tAI plotted against CAI quintiles.* ***A.*** *Global CAI.* ***B.*** *Regional CAI.*

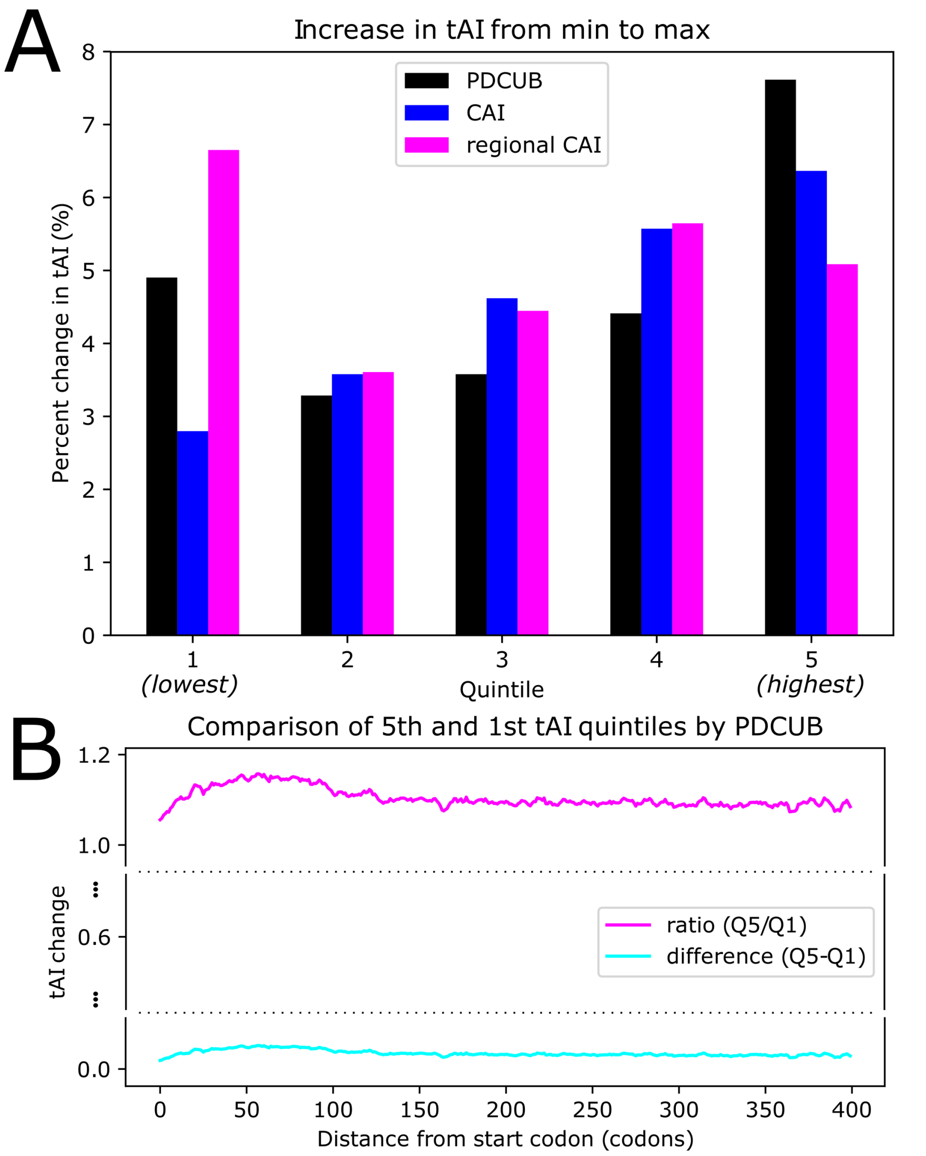


***Supplementary Figure 10:*** *Comparison of increase in tAI by quintile.* ***A.*** *Side-by-side percent increase in tAI from lowest point to highest point for each PDCUB, CAI, and regional CAI quintile.* ***B.*** *Ratio of, and difference between, tAI for fifth and first PDCUB quintiles.*

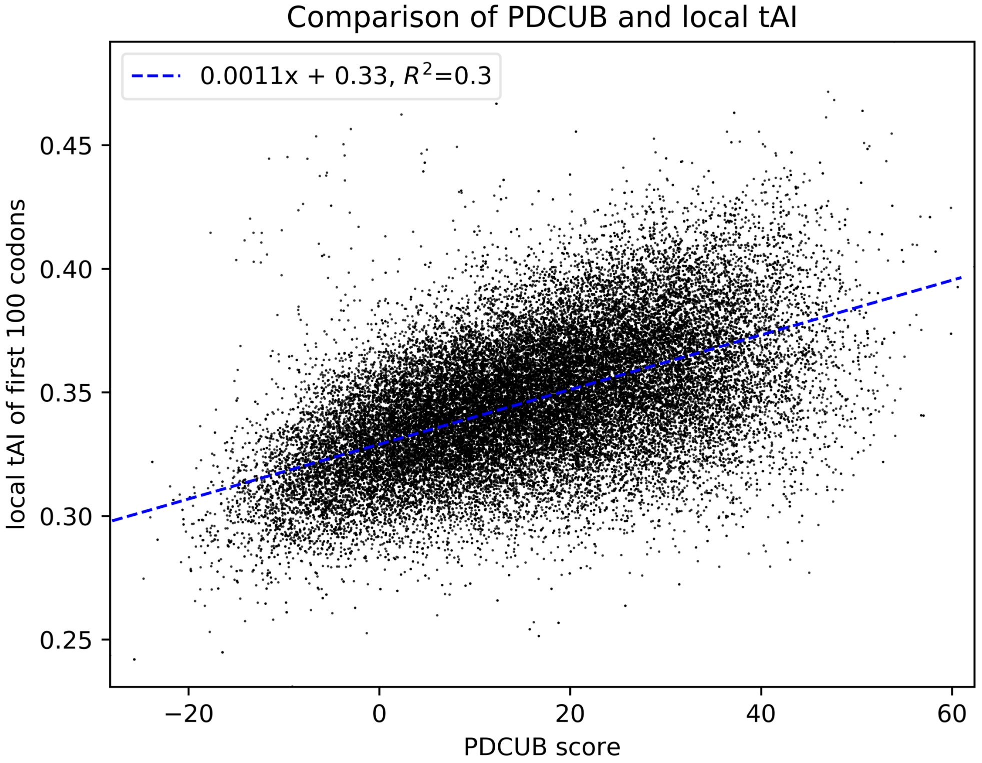


***<u>Supplementary Figure 11:</u>*** *Scatter plot of local tAI for the first 100 codons of all coding transcripts, plotted against the PDCUB score of the same transcript. Each dot corresponds to a transcript. The dashed blue line is the best linear fit to the scatter data.*

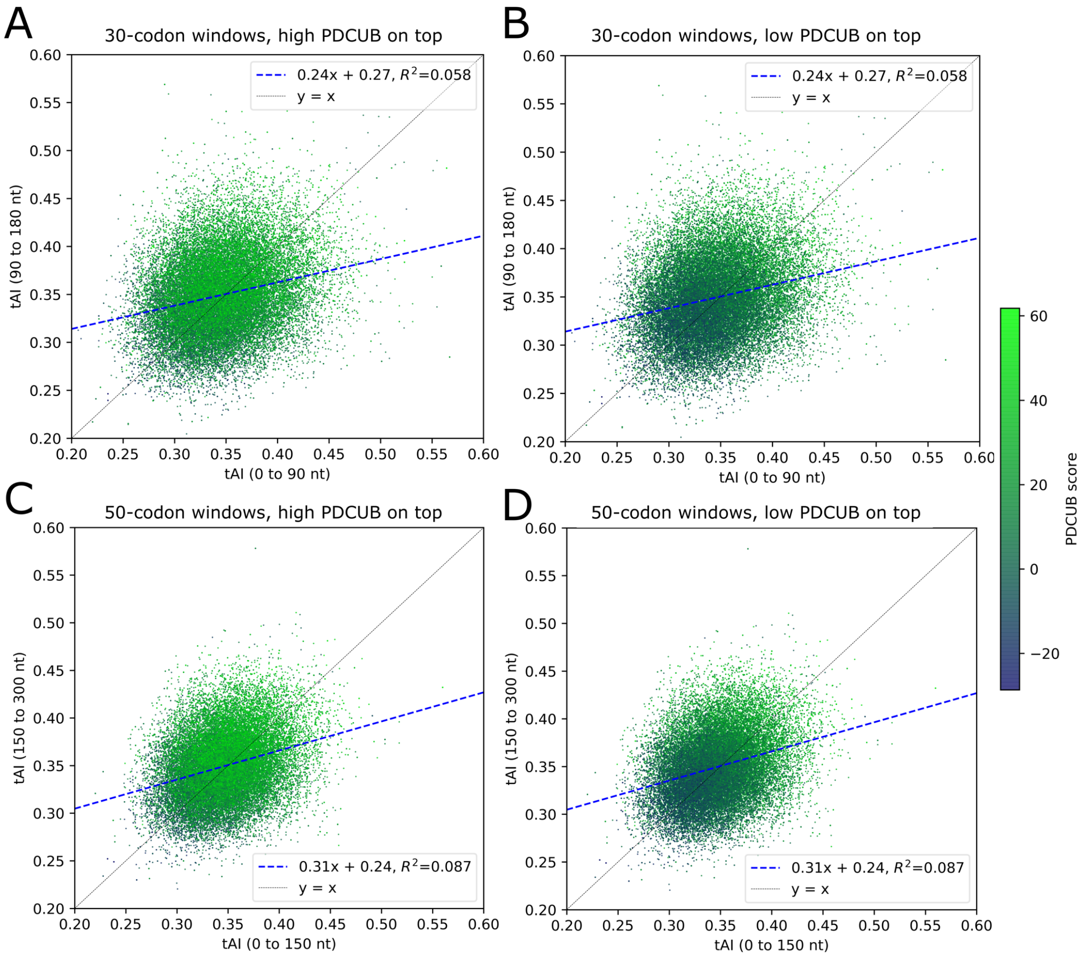


***Supplementary Figure 12:*** *Scatterplots of relative local tAI for first and second bins in initial regions of coding transcripts. Each point corresponds to a single CDS, while the colormap indicates PDCUB score.* ***A.*** *Second bin of 30 codons plotted against first bin of 30 codons, with highest-PDCUB transcripts on top.* ***B.*** *Same as A, but with lowest-PDCUB transcripts on top.* ***C.*** *Second bin of 50 codons plotted against first bin of 50 codons, with highest-PDCUB transcripts on top.* ***D.*** *Same as C, but with lowest-PDCUB transcripts on top.*

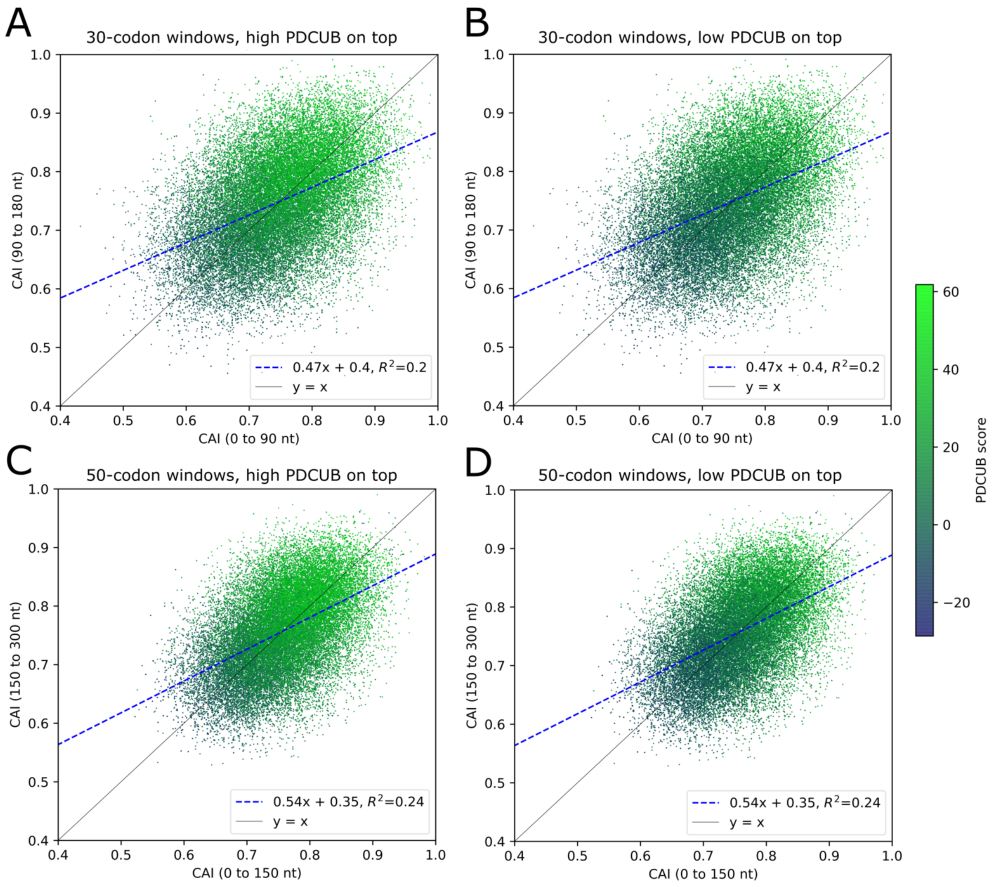


***Supplementary Figure 13:*** *Scatterplots of relative regional CAI (global RA) for first and second bins in initial regions of coding transcripts. Each point corresponds to a single CDS, while the colormap indicates PDCUB score. Low-scoring PDCUB sequences show the greatest deviation from y=x, while high-PDCUB transcripts correspond to a high regional CAI for both regions of the CDS.* ***A.*** *Second bin of 30 codons plotted against first bin of 30 codons, with highest-PDCUB transcripts on top.* ***B.*** *Same as A, but with lowest-PDCUB transcripts on top.* ***C.*** *Second bin of 50 codons plotted against first bin of 50 codons, with highest-PDCUB transcripts on top.* ***D.*** *Same as C, but with lowest-PDCUB transcripts on top.*

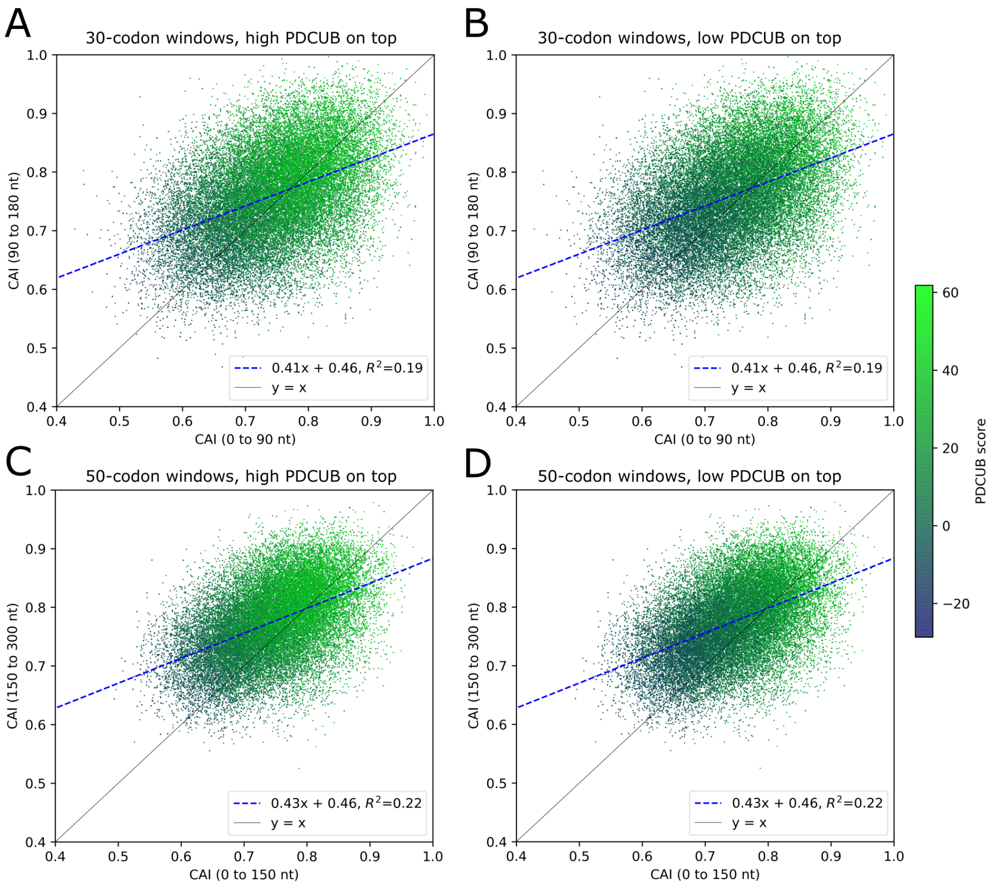


***Supplementary Figure 14:*** *Scatterplots of relative regional CAI (regional RA) for first and second bins in initial regions of coding transcripts. Each point corresponds to a single CDS, while the colormap indicates PDCUB score. Low-scoring PDCUB sequences show the greatest deviation from y=x, while high-PDCUB transcripts correspond to a high regional CAI for both regions of the CDS.* ***A.*** *Second bin of 30 codons plotted against first bin of 30 codons, with highest-PDCUB transcripts on top.* ***B.*** *Same as A, but with lowest-PDCUB transcripts on top.* ***C.*** *Second bin of 50 codons plotted against first bin of 50 codons, with highest-PDCUB transcripts on top.* ***D.*** *Same as C, but with lowest-PDCUB transcripts on top.*

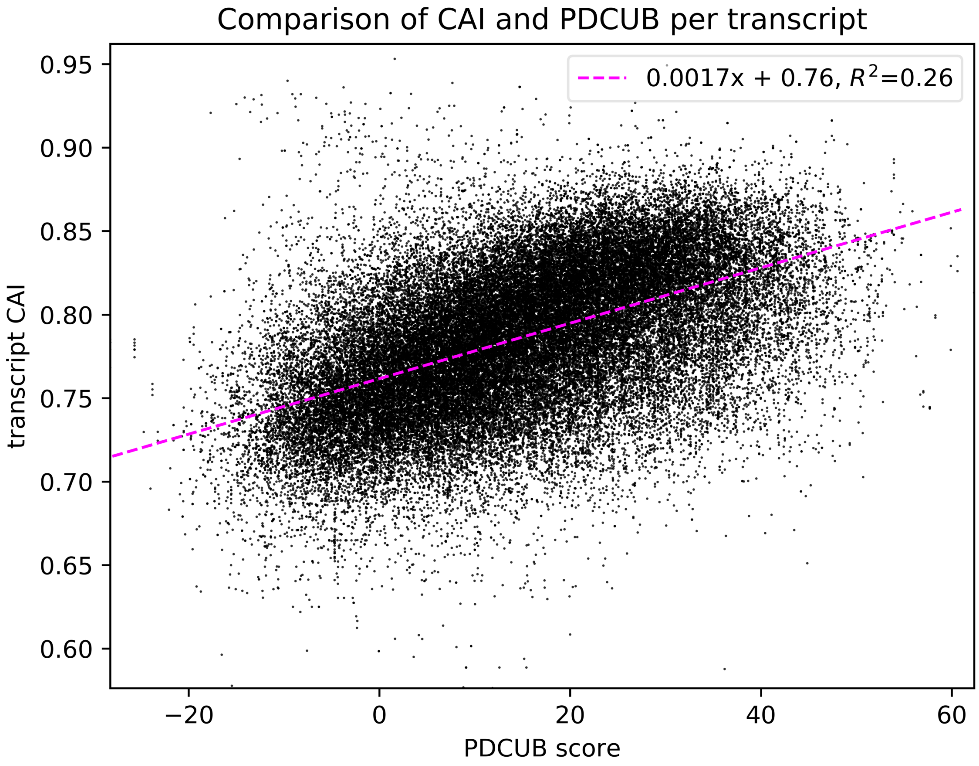


***Supplementary Figure 15:*** *Scatter plot of CAI for each coding transcript plotted against the PDCUB score of the same transcript. Each dot corresponds to a transcript. The dashed magenta line is the best linear fit to the scatter data.*

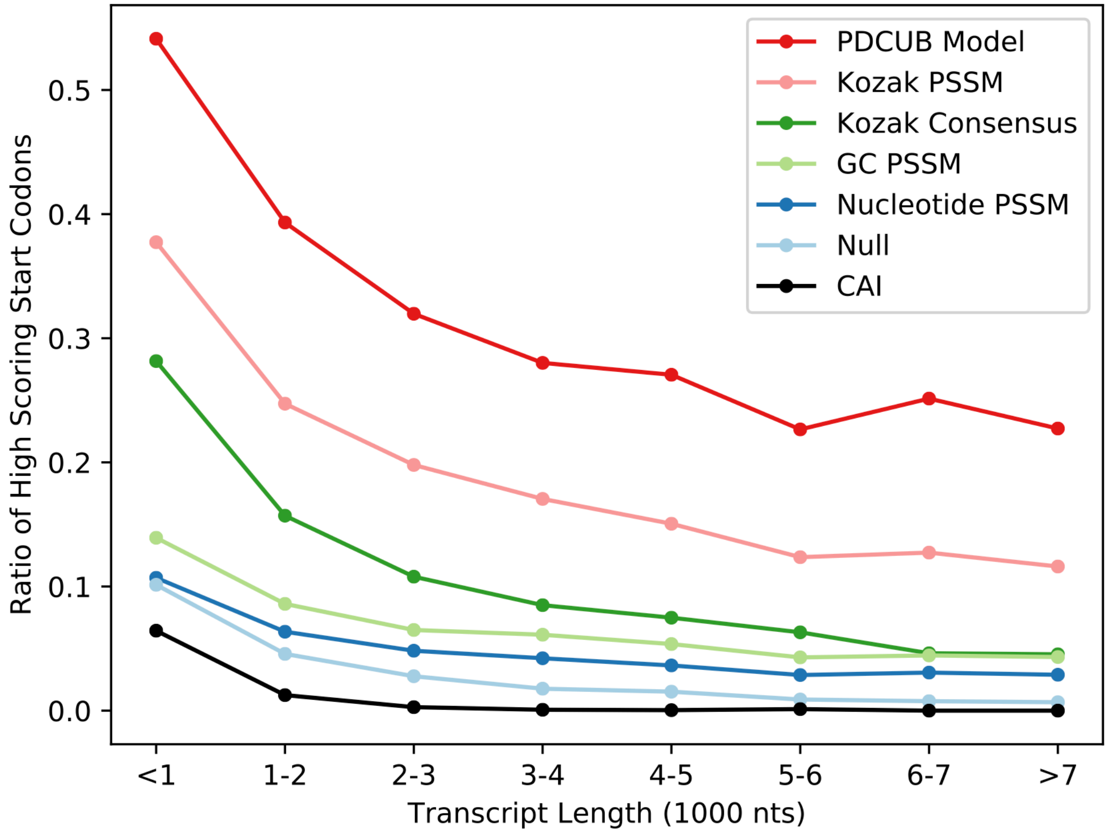


***Supplementary Figure 16****: The rate of start codon identification compared for different models, including CAI. The curves show what fraction of start codons are identified per transcript, comparing the PDCUB score with other models, similarly as Figure 3A, but with CAI computed for the first 100 codons also compared.*

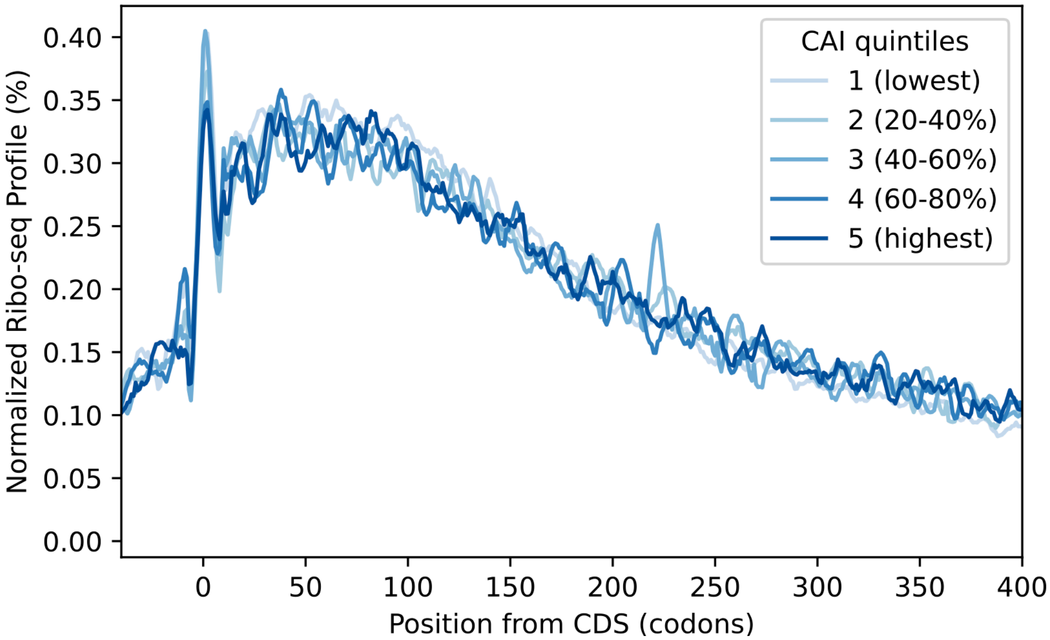


***<u>Supplementary Figure 27</u>****: The shape of ribosomal occupancy, shown as the normalized Ribo-seq profile centered at the start codon at position 0 for each quintile defined by CAI. Read positions are A-shifted by 15-nt, and smoothed using a Savitzky-Golay filter.*
